# Glycan-dependent two-step cell adhesion mechanism of Tc toxins

**DOI:** 10.1101/857730

**Authors:** Daniel Roderer, Felix Bröcker, Oleg Sitsel, Paulina Kaplonek, Franziska Leidreiter, Peter H. Seeberger, Stefan Raunser

**Affiliations:** Department of Structural Biochemistry, Max Planck Institute of Molecular Physiology, Otto-Hahn-Str. 11, 44227 Dortmund, Germany; Department of Biomolecular Systems, Max Planck Institute of Colloids and Interfaces, Am Mühlenberg 1, 14476 Potsdam, Germany; Vaxxilon Deutschland GmbH, Magnusstr. 11, 12489 Berlin, Germany; Department of Biomolecular Mechanisms, Max Planck Institute for Medical Research, Jahnstraße 29, 69120 Heidelberg, Germany

## Abstract

Toxin complex (Tc) toxins are virulence factors widespread in insect and human bacterial pathogens. Tcs are composed of three subunits: TcA, TcB and TcC. TcA facilitates receptor-toxin interaction and membrane permeation, TcB and TcC form a toxin-encapsulating cocoon. While the mechanisms of holotoxin assembly and prepore-to-pore transition have been well-described, little is known about receptor binding and cellular uptake of Tcs. Here, we identify two classes of glycans, heparins/heparan sulfates and Lewis antigens, that act as receptors for different TcAs from insect- and human pathogenic bacteria. Glycan array screening and electron cryo microscopy (cryo-EM) structures reveal that all tested TcAs bind unexpectedly with their *α*-helical part of the shell domain to negatively charged heparins. In addition, TcdA1 from the insect-pathogen *Photorhabdus luminescens* binds to Lewis antigens with micromolar affinity. A cryo-EM structure of the TcdA1-Lewis X complex reveals that the glycan interacts with the receptor-binding domain D of the toxin. Our results suggest a two-step association mechanism of Tc toxins involving glycans on the surface of host cells.

## Introduction

A widespread type of toxin in insect and human bacterial pathogens is the heterotrimeric toxin complex (Tc)^1, 2^. Tc toxins were originally discovered in the insect pathogen *Photorhabdus luminescens*, which lives in symbiosis with soil nematodes^3^. Later, gene loci of Tc toxins were also found, amongst others, in *Xenorhabdus nematophila*^4, 5^, in the facultative human-pathogen *Photorhabdus asymbiotica*^6^ and different species of *Yersinia*, which are both human and insect pathogens^7, 8^.

Tc toxins consist of three components: TcA, TcB and TcC. The ∼ 1.4 MDa TcA is a homo-pentameric bell-shaped molecule that mediates target cell association, membrane penetration and toxin translocation^9^. TcA is made up of a central, pre-formed channel that is enclosed by an outer shell composed of a structurally conserved *α*-helical domain decorated with variable receptor-binding domains (RBDs). The shell and channel are connected by a stretched linker^10^. TcB and TcC together form a ∼ 250 kDa cocoon that encapsulates the toxic component, the ∼ 30 kDa C-terminal hypervariable region (HVR) of TcC, which is autoproteolytically cleaved and resides inside the cocoon^10, 11^.

Binding of TcB-TcC to TcA involves a conformational transition of the TcA-binding domain of TcB, which is a distorted six-bladed *β*-propeller that closes the cocoon at the bottom. Following the contact of TcA and TcB, two blades of the *β*-propeller unfold and refold, resulting in the opening of the cocoon. As a consequence, the toxic enzyme enters the translocation channel of TcA^12^.

After specific binding to receptors on the surface of the target cell, the holotoxin is endocytosed^13^. Acidification in the late endosome triggers the opening of the shell and the compaction of the stretched linker drives the release of the channel from the shell and its entry into the target cell membrane. In the membrane the tip of the channel opens and enables the translocation of the actual toxin^14, 15^.

While the mechanisms of holotoxin assembly and prepore-to-pore transition have been well-described (reviewed in ^16^), little is known regarding the receptor binding and cellular uptake of Tc toxins. In the case of YenTcA, the TcA from the insect pathogen *Yersinia entomophaga*, two chitinases form a complex with the toxin^17^. It was proposed that the chitinases cause the degradation of the peritrophic membrane within the midgut of insects to facilitate toxin entry^18^. The association of YenTcA with target cells potentially occurs via glycan structures, which were recently identified in a binding screen to associate with both chitinases and directly with YenTc^19^.

Various other toxins have been shown to bind to glycans on target cells, such as botulinum neurotoxin, which binds to *N*-glycosylated SV2^20^, cholera toxin, which binds to ganglioside GM1^21^, and typhoid toxin from *Salmonella typhi*, which binds to multiantennal glycans^22^. The structural similarity of the shell-enclosing RBDs of various TcAs to sialidase^10, 23^ suggests that glycans could be also the cellular receptors in the case of TcAs that do not associate with chitinases.

Here, we used a synthetic glycan microarray^24^ to specifically screen for glycans as possible receptors for Tc toxins from several insect and human pathogens (*P. luminescens* TcdA1 (Pl-TcdA1), *X. nematophila* XptA1 (Xn-XptA1), *Morganella morganii* TcdA4 (Mm-TcdA4), *Y. pseudotuberculosis* TcaATcaB (Yp-TcaATcaB)). All tested TcAs interact with heparins/heparan sulfates (HS). The cryo-EM structure of Mm-TcdA4 in complex with heparin reveals that the glycan interacts directly with the *α*-helical part of the shell domain. In addition, Pl-TcdA1 interacts with several Lewis antigens, which bind to the RBD D as revealed by the cryo-EM structure of the Pl-TcdA1 in complex with BSA-Lewis X.

## Results

### Complex glycans on the cell surface mediate association of TcA

When intoxicating different cell types with Pl-TcdA1, we found that HEK 293 GnTI^-^ cells are less susceptible to the toxin than HEK 293T cells (Supplementary Figure 1a,b). While HEK 293T cells readily bind and accumulate fluorescently labeled Pl-TcdA1 after one to four hours of incubation time, HEK 293 GnTI^-^ cells bind considerably less fluorescent Pl-TcdA1 (Figure 1a,b). The major difference between the two cell types is that HEK 293 GnTI^-^ cells do not have *N*-acetyl-glucosaminyltransferase I, therefore lack complex *N*-glycans^25^. The lack of mature N-linked glycans on the plasma membrane of HEK 293 GnTI^-^ cells is probably responsible for the decreased interaction with Pl-TcdA1, suggesting that *N*-linked glycans play a major role in the specific toxin binding to the host membrane.

**Figure 1:**
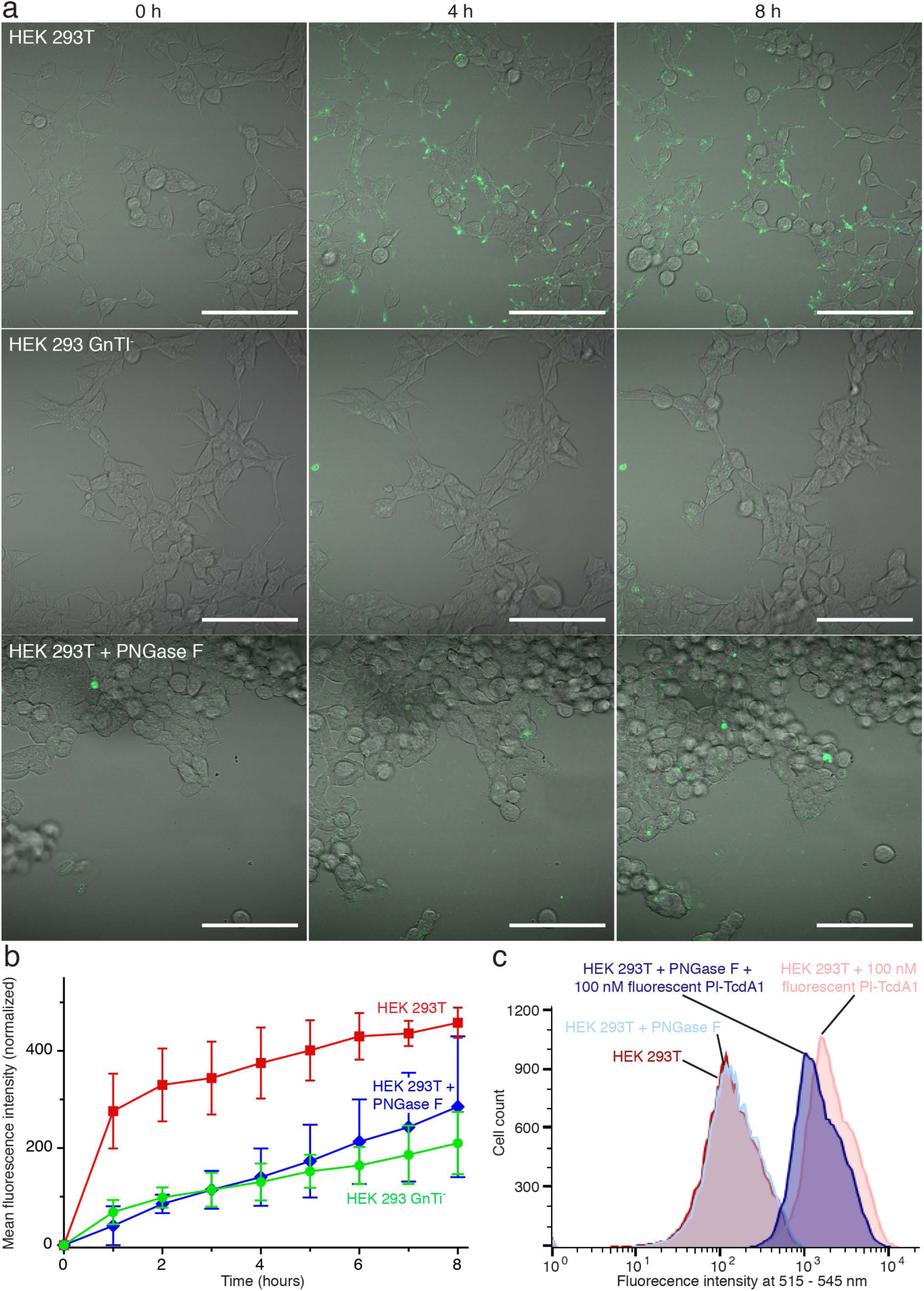
The presence of complex glycans on target cells enhances binding of Pl-TcdA1. a: Time course confocal imaging demonstrates that wild type HEK 293T cells bind Pl-TcdA1 (AlexaFluor488 labelled, green) much more readily than their glycosylation-deficient HEK 293 GnTI^-^ counterparts and PNGase F-treated HEK 293T cells. Scale bar, 100 µm; fluorescence channels are equally thresholded. b: Quantification of AlexaFluor488 labelled Pl-TcdA1 binding to wild type HEK 293T, PNGase F-treated HEK 293T and HEK 293 GnTI^-^ cells based on time course confocal microscopy data. Three biological replicates for each cell type were analyzed; error bars represent standard deviation. c: Flow cytometry of HEK 293T exposed to AlexaFluor488 labelled Pl-TcdA1 after deglycosylation of cell surface by PNGase F. Histograms of PNGase F-treated and untreated cells, both with and without 100 nM Pl-TcdA1 are shown in comparison. The gating strategy and cell numbers are shown in Supplementary data file 1.

To confirm that this is indeed the case, we treated HEK 293T cells with PNGase F to completely remove *N*-linked glycans on the plasma membrane^26^. Pl-TcdA1 binding to these cells was tremendously decreased supporting our conclusion that *N*-linked glycans are important for the binding of the toxin to host cells (Figure 1c, Supplementary Figure 1b).

### Pl-TcdA1 binds to Lewis antigens

In order to identify possible glycans that are responsible for the specific binding of Tc toxins to host cells, we first applied a synthetic glycan microarray and screened TcAs from different organisms, namely Pl-TcdA1, Xn-XptA1, Mm-TcdA4, and Yp-TcaATcaB. We identified several Lewis oligosaccharides that interacted with Pl-TcdA1 (Figure 2a,b). Especially the trisaccharide Lewis X and the tetrasaccharides Lewis Y and sialyl-Lewis X interacted strongly with Pl-TcdA1 (Figure 2b,c, Supplementary Figure 2a). Interestingly, none of the other TcAs showed a significant binding to the microarray even at higher protein concentrations, indicating that this glycan-toxin interaction is specific for Pl-TcdA1.

**Figure 2:**
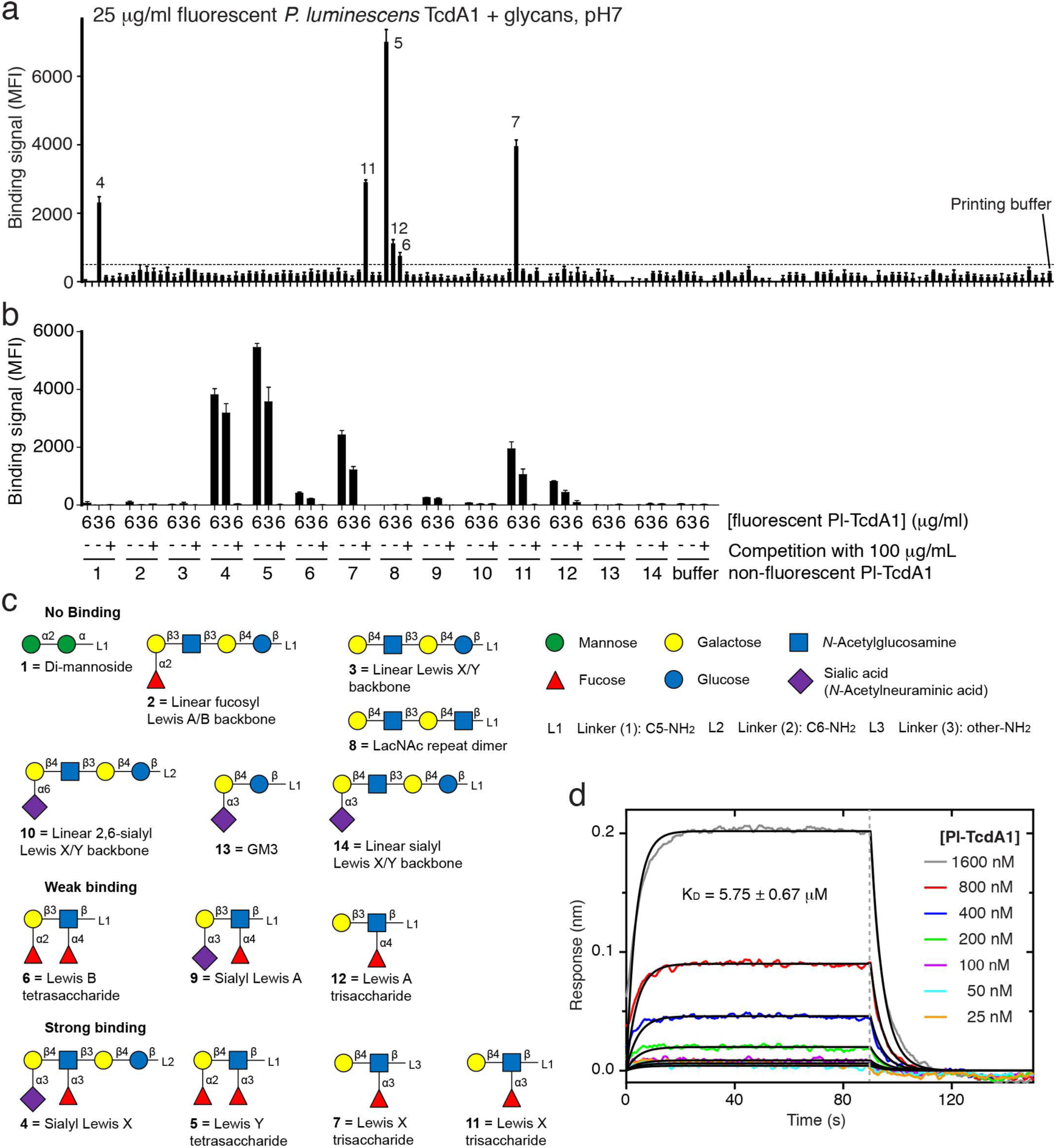
Interaction of Pl-TcdA1 with various Lewis antigens. a: Glycan microarray showing the interaction of fluorescently labeled Pl-TcdA1 (fl-TcdA1) with various glycans. Binding signals are shown as local background-subtracted mean fluorescence intensity (MFI) values to 141 synthetic glycans. MFI values are indicated as bars (mean ± standard deviation). An arbitrary threshold of MFI=500 is indicated as dashed line. b: Focused microarray with selected glycans from (a). 6 or 3 μg/mL of fl-TcdA1 were applied to 14 different glycans (indicated below the bar diagrams, spotted at 0.1 mM) in the absence or presence of an excess of unlabeled Pl-TcdA1. c: Schematic representation of glycans used on the focused microarray, grouped by binding of Pl-TcdA1 (no binding, weak binding and strong binding). The individual monosaccharide moieties are shown on the top right. d: BLI sensorgrams of TcdA1 interacting with immobilized BSA-Lewis X. TcA pentamer concentrations were 25 – 1600 nM. A global fit according to a 1:1 binding model was applied (black curves), resulting in a K_D_ of 5.75±0.67 μM, k_on_ of 3.81±0.44*×*10^4^ M^-1^ s^-1^, and k_off_ of 2.19±0.04*×*10^-1^ s^-1^. Association and dissociation phases are separated by a grey dotted line.

The interaction with Lewis oligosaccharides is unexpected since these *α*1,3-fucosylated glycans are normally found on the surface of human red blood cells^27^ and not in insects. Therefore, it is unlikely that they represent the natural receptor of Pl-TcdA1, which is a Tc toxin from an insect pathogenic bacterium. Although Lewis antigens have not been described for insects, similar oligosaccharides with *α*1,3-fucosylated Lewis-like antennae have been reported to be present in glycoproteins of the lepidopterans *Trichoplusia ni* and *Lymantria dispar*^28^. Interestingly, *Lepidoptera* are typical hosts of nematodes living in symbiosis with *P. luminescens*^3^ and are therefore the natural target of Tc toxins. We therefore believe that the interaction between Pl-TcdA1 and Lewis antigens is representative for the binding of the toxin to similar glycan receptors on the insect host membrane.

Protein-glycan interactions are generally weak and multivalent interactions are in many cases necessary to achieve a significant increase in receptor binding affinity^29^. Since TcA is a homopentameric complex, high-affinity receptor binding is likely also achieved by multivalency. To determine the binding kinetics of Lewis X antigens to TcA pentamers, we conjugated multiple oligosaccharides to BSA as carrier protein (Supplementary Figure 2b-d) and quantified the protein-oligosaccharide interaction using biolayer interferometry (BLI). This approach mimics the high surface density of glycans on a cell surface and thereby ensures that all Lewis X binding sites on a TcA pentamer can be occupied.

We determined a dissociation constant (K_D_) of 5.8 ± 0.7 μM (Figure 2d) for the complex of Pl-TcdA1 and the BSA-Lewis X glycoconjugate. The relatively high on-and off-rates of 3.8 ± 0.4 *×*10^4^ M^-1^ s^-1^ and 0.22 ± 0.003 s^-1^, respectively, indicate a highly dynamic binding equilibrium between Lewis X and Pl-TcdA1. This probably provides the toxin with a certain mobility on the cell surface and allows it to laterally move to regions with high receptor density.

### Structure of the Pl-TcdA1-BSA-Lewis X complex

To elucidate how Lewis X interacts with Pl-TcdA1, we planned to determine the structure of the Pl-TcdA1-BSA-Lewis X complex by single particle cryo-EM. However, since the affinity of Pl-TcdA1 to BSA-Lewis X is low, high amounts of BSA-Lewis X are required to allow for complex formation which is incompatible with single particle cryo-EM. Therefore, we stabilized the Pl-TcdA1-BSA-Lewis X complexes that were formed using a large excess of BSA-Lewis X by crosslinking with glutaraldehyde and removed unbound BSA-Lewis X. The resulting complexes were suitable for single particle cryo-EM (Supplementary Figure 3a). Analyzing the single particles by two-dimensional clustering and sorting in SPHIRE^30^ revealed up to three additional densities corresponding to BSA-Lewis X at the periphery of Pl-TcdA1, indicating that the glycan interacts with one of the RBDs of the toxin (Figure 3a, Supplementary Figure 3b). Since not all five Lewis X binding sites of the Pl-TcdA1 pentamer are occupied and the number of BSA-Lewis X molecules per complex varies, we combined three-dimensional classification with symmetry expansion (Methods, Supplementary Figure 4) to obtain a three-dimensional reconstruction of the Pl-TcdA1-BSA-Lewis X complex, in which one BSA-Lewis X is bound to one Pl-TcdA1 pentamer (Figure 3b).

**Figure 3:**
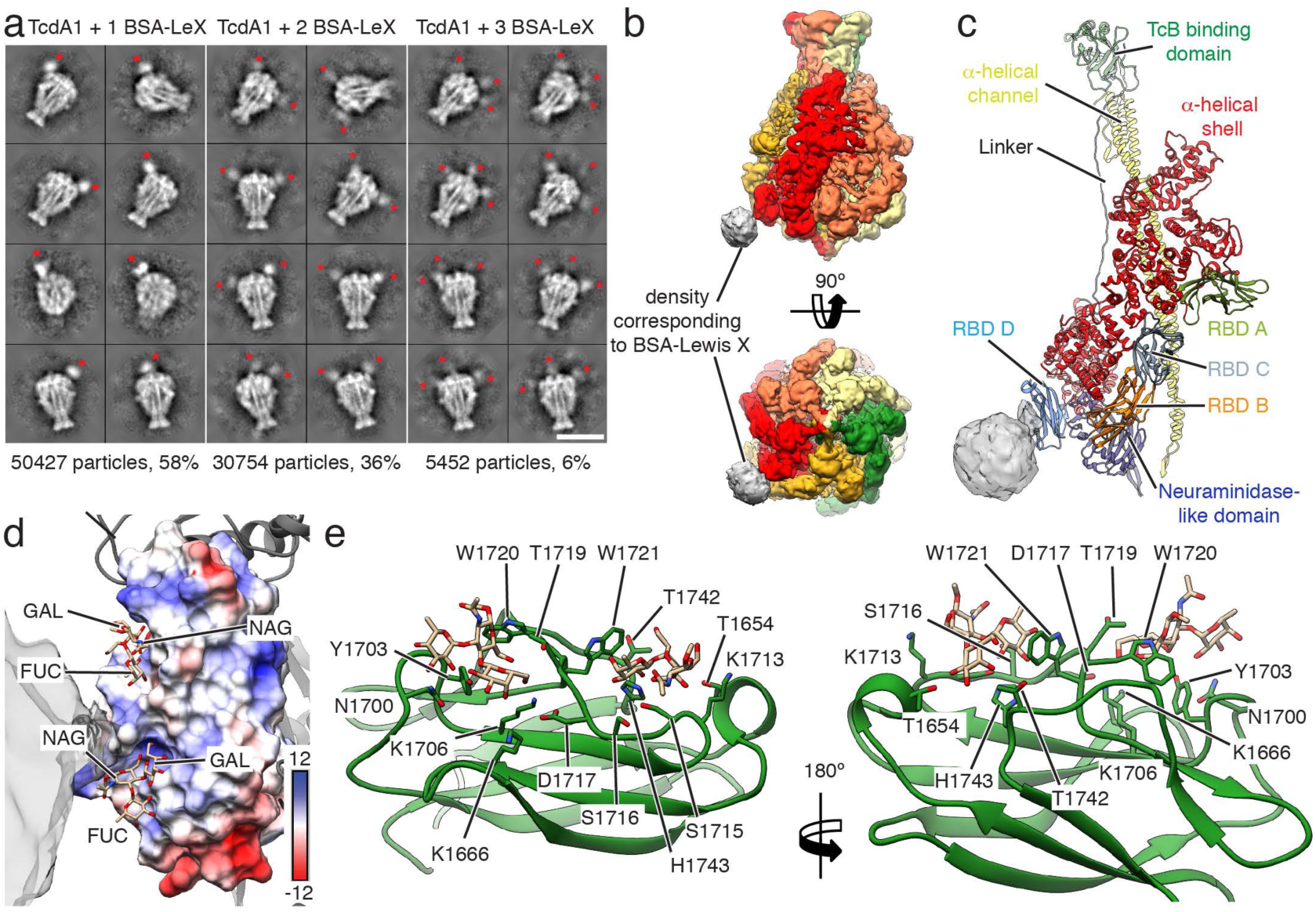
Structure of Pl-TcdA1 with BSA-Lewis X. a: Representative reference-free 2D class averages of cryo-EM images of Pl-TcdA1 with crosslinked BSA-Lewis X obtained by ISAC and resampled to the original pixel size. Scale bar, 20 nm. b: Cryo-EM map of Pl-TcdA1 with BSA-Lewis X. The protomers are colored individually. Density corresponding to BSA-Lewis X (gray) is shown at lower binarization threshold. C: Model of one protomer of Pl-TcdA1 together with the cryo-EM density of BSA-Lewis X (gray) at low binarization threshold. The domains of Pl-TcdA1 are colored individually. BSA-Lewis X is attached to RBD D. d: Surface representation of RBD-D, colored according to the Coulomb potential (kcal mol^-1^ e^-1^) at pH 7.0. Two docked conformations of Lewis X are shown as stick representations. The docking score is −4.5 kcal/mol for both conformations. The cryo-EM density map corresponding to BSA-Lewis X is transparent gray. e: Detailed view of RBD-D (green) with two docked Lewis X molecules. Side chains that contact the docked molecules are highlighted.

Despite the heterogeneity of the data set and the flexibility of the bound BSA-Lewis X complex, the reconstruction reached an average resolution of 5 Å (Supplementary Figure 3c-e). This enabled us to accurately fit the previously determined cryo-EM structure of Pl-TcdA1 into the toxin density^23^ (Figure 3c). Although the density corresponding to BSA-Lewis X is less well resolved and therefore the glycan or BSA cannot be fitted, its position on the toxin unambiguously demonstrates that it binds to RBD D (Figure 3b,c, Supplementary Movie 1).

Molecular docking of the Lewis X trisaccharide on RBD D revealed two prominent potential binding sites (Figure 3d, e, Supplementary Movie 1). Separated by two tryptophan residues (W1720 and W1721), both binding sites form positively charged grooves with many polar residues (predominantly Thr, Ser and Lys), allowing hydrogen bond mediated interactions with the glycan.

### TcAs of insect-and human-pathogenic bacteria bind to various heparins

To find out whether also heparins/heparan sulfates (HS) interact with the four different TcAs, we performed a glycan screen comprising different heparins/HS oligosaccharides. Heparins and HS are the most structurally complex glycosaminoglycans and are a major component of the extracellular space^31^. They are composed of disaccharide repeating units of D-glucosamine (GlcN) and either l-iduronic acid (IdoA) or D-glucuronic acid (GlcA) linked by α1-4 and β1-4 glycosidic bonds, respectively. Sulfation can occur at positions 2, 3 and 6 of GlcN as well as at position 2 of IdoA/GlcA. Heparin is primarily expressed in mast cells, while HS is ubiquitous on cell surfaces and in the extracellular matrix. In heparin, uronic acid residues are 90% IdoA and 10% its C5 epimer GlcA. The prototypical heparin disaccharide contains three sulfate groups (2.7 sulfates per disaccharide on average in the polymer). HS chains are typically more heterogeneous than those of heparin, are richer in *N*-acetyl D-glucosamine (GlcNAc) and GlcA and contain fewer O-sulfates (one per disaccharide on average in the polymer). We found that all tested TcAs interact with heparin or HS (Figure 4a). The binding signal was especially strong for Xn-XptA1 and Mm-TcdA4. The controls (wheat germ agglutinin and BSA) almost did not bind to the microarray (Figure 4a), indicating that the heparin/HS-toxin interaction is indeed specific. While the chain length of the strong binding heparins/HSs varies and seems therefore not to be crucial for the interaction (Figure 4b), we observed that all of them had at least two negative charges per monosaccharide group, suggesting that the binding of TcA to heparins/HS is facilitated by electrostatic interactions.

**Figure 4:**
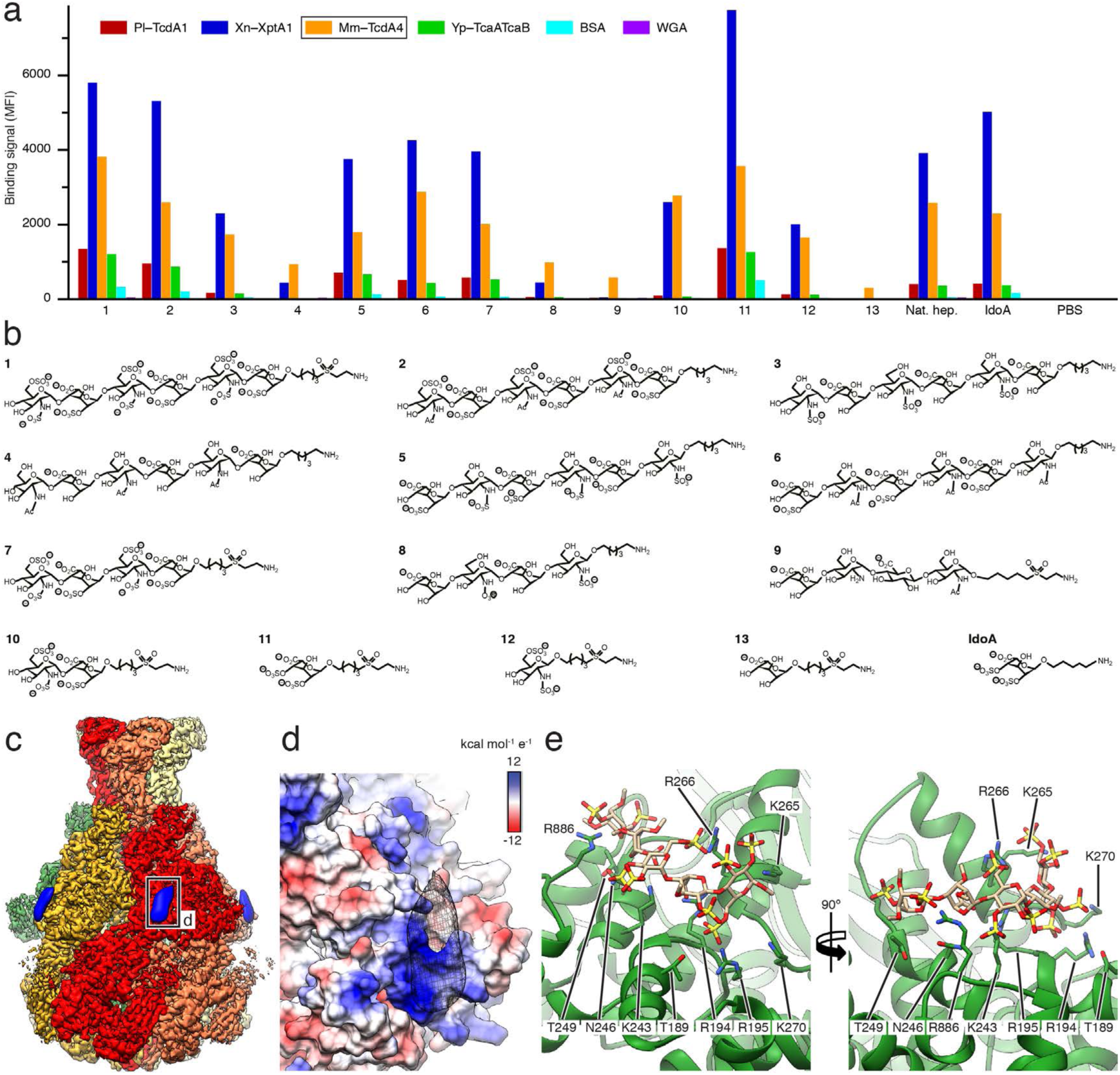
Interaction of TcAs with various heparin oligosaccharides and structure of Mm-TcdA4 in complex with heparin. a: Glycan microarray showing the interaction of the indicated TcAs (0.25 μg/mL) with selected heparin oligosaccharides, the natural heparin mixture from porcine intestinal mucosa (Nat. hep.), and the heparin analogue IdoA-2,4-disulfate(a)linker (IdoA). As controls bovine serum albumin (BSA) and wheat germ agglutinin (WGA) were used. As an additional negative control phosphate-buffered saline (PBS) was spotted on the array. The full heparin microarray is shown in Supplementary Figure 5a. b: Structures of heparins 1-13 and IdoA. The numbers correspond to (a). c: Cryo-EM structure of Mm-TcdA4 (highlighted in (a)) in complex with natural heparin. The protomers of Mm-TcdA4 are colored individually. Additional density not corresponding to the toxin is filtered to 15 Å and shown in blue. d: Surface representation of the heparin binding site of the shell domain of Mm-TcdA4, colored according to the Coulomb potential (kcal mol^-1^ e^-1^) at pH 7.0. The additional density corresponding to heparin is shown in black mesh representation. e: The interface region of the Mm-TcdA4 shell domain (green) with side chains that contact a docked heparin pentasaccharide (sand colored). The docking score is −6.2 kcal/mol. All residues belong to the *α*-helical part of the shell.

### Structure of the Mm-TcdA4-heparin complex

We chose Mm-TcdA4 from the opportunistic human pathogen *Morganella morganii* to investigate the heparin/HS-toxin interaction in more detail. First, we performed biolayer interferometry (BLI) to determine the affinity of Mm-TcdA4 to heparin/HS. Since synthetic heparins/HSs are difficult to produce in large amounts needed for biophysical and structural studies, we decided to work with natural ∼15 kDa porcine intestinal mucosa heparin instead, albeit its heterogeneity. The small apparent average dissociation constant (K_D_) of the heparin mixture (0.23 ± 0.01 nM) clearly demonstrates a high binding affinity of the glycans to Mm-TcdA4 (Supplementary Figure 5).

To identify which domains of Mm-TcdA4 are involved in heparin/HS binding, we determined the structure of Mm-TcdA4 in complex with porcine intestinal mucosa heparin using single particle cryo-EM (Supplementary Figure 6, Supplementary Movie 2). The 3.2 Å structure is very similar to the previously obtained structure of Mm-TcdA4^23^ (Supplementary Figure 7a). However, we identified additional densities at the periphery of the shell domain of each protomer (Figure 4c, Supplementary Figure 7b,c, Supplementary Movie 2). The density occupies a cleft that is predominantly formed by positively charged residues (R194, R195 K243, K265, R266 and R886), representing the ideal binding site for negatively charged heparins (Figure 4d). This clearly shows that the heparins do not interact with an RBD, but directly with the *α*-helical domain of the shell. Although the resolution of the density does not allow to build an atomic model in this region, we flexibly fitted several heparin oligosaccharides that resemble natural heparin into the density. Indeed, most of them fit well and their negatively charged residues are involved in many potential electrostatic interactions at the interface (Figure 4e, Supplementary Figure 7d).

## Discussion

In the current work, we demonstrate that Tc toxins specifically interact with glycans on the surface of host cells. We identified two specific types of glycans that act as receptors for Tc toxin binding: heparins/HSs and Lewis antigens. While heparins bind to a specific groove of the *α*-helical shell domain of all tested TcAs, Lewis antigens only interact with the RBD D of Pl-TcdA1.

The heparin/HS binding site in the *α*-helical shell domain is 10 nm away from the membrane if the Tc toxin is in the prepore conformation and attaches perpendicular to the membrane. HS proteoglycans form long polymers on cell surfaces that extend 20–150 nm from the plasma membrane^32, 33^. Thus, they can easily bind to the toxin even before it reaches the membrane surface. We propose that Tc toxins bind first to HS, which brings them closer to the cell surface and thereby in contact with other surface receptor molecules. HS also bind to viruses (e.g. baculovirus^34^) and cationic cell-penetrating peptides^35^ that act as their respective receptors for cell entry. *Staphylococcus aureus* beta-toxin uses the heparan sulfate carrying protein syndecan-1 as a cellular receptor^36^.

Lewis antigens on the other hand are shorter glycans than the glycosaminoglycans. Thus, the toxins have to be closer to the membrane in order to interact with them. We therefore propose that Pl-TcdA1 binds in a second step to Lewis antigens.

Because the five identical RBD Ds which interact with Lewis antigens are symmetrically arranged at the bottom of the Pl-TcdA1 shell, the binding affinity is highest when the toxin is oriented perpendicular to the membrane. This puts the toxin in the ideal position for membrane penetration. Our data show that the interaction of Pl-TcdA1 with BSA-Lewis X is highly dynamic, which could allow for the toxin to move laterally on the membrane surface. Such an effect has been previously described for bacteria and viruses that interact via lectins with the host cell surface^32^.

Lewis antigens are normally found on the surface of human red blood cells^27^ and have been identified as receptors for toxins of human pathogens and viruses, such as cholera toxin^37^ or norovirus^38^. However, Lewis-like oligosaccharides with core *α*1,3-fucosylation also have been identified in various insect species, making them likely receptors for Tc toxins from entomopathogenic bacteria.

In a previous glycan array screen with the Tc toxin Yen-Tc from the insect pathogen *Yersinia entomophaga*, 72 out of 432 glycans were identified as potential toxin receptors, amongst them monosaccharides, fucosylated glycans, mannose derivatives, glucose derivatives, sialylated derivatives, glycosaminoglycans, complex *N*-glycans and derivatives with terminal GlcNAc, GalNAc or galactose^19^. In contrast, Xn-XptA1, Mm-TcdA4 and Yp-TcaATcaB only interacted strongly with heparins/HS (11 out of 36 heparins/HS on the microarray). Pl-TcdA1 additionally bound to core-fucosylated Lewis antigens (6 out of 141 glycans on the microarray). This indicates that the chitinase-associated Yen-Tc is clearly less selective in terms of glycan binding than the TcAs that we tested here.

What about the other RBDs? Their structure is similar to RBD D, suggesting that they would also interact with glycans that we have not identified so far. There are three possible explanations why these interactions may have not been detected by our microarray screen. First, the distance between the glycans on the array surface and the RBDs in the toxins differ from the situation on cell surfaces, where the glycans are mostly bound to proteins. Therefore, possible glycan binding especially to RBDs that locate distally to the membrane might be undetectable, especially when considering the effect of multivalency in the pentameric TcA. Second, our glycan screen covers only a selection of glycans that exist on target cells. For example multi-antennal glycans, which have been described to function as high-affinity receptors for typhoid toxin^22^, were not present in the array. Third, it could well be that several of the RBDs do not bind to glycans on the cell surface, but directly to proteins.

The removal of *N*-linked glycans on HEK 293T cells or the absence of complex glycans on HEK 293 GnTi^-^ cells reduced the number of Pl-TcdA1 toxins bound to the cell surface when compared to untreated HEK 293T cells, but it did not completely prevent toxin binding. This indicates that the toxin does not only bind to *N*-linked glycans, but probably also interacts with *O*-linked glycans and/or directly with proteins or lipids in the host cell membrane.

Taken together, our data allow us to propose the following model for Tc toxin binding to cells (Figure 5). Initially, Tc toxins bind to long heparan sulfate glycosaminoglycans to bring them into close contact to other surface receptor molecules. In a second step, the toxin interacts with these receptors to orient the toxin in an ideal position for membrane penetration. Next, the toxin undergoes endocytosis. Acidification in the late endosome triggers then the prepore-to-pore transition and membrane permeation of TcA (Figure 5).

**Figure 5:**
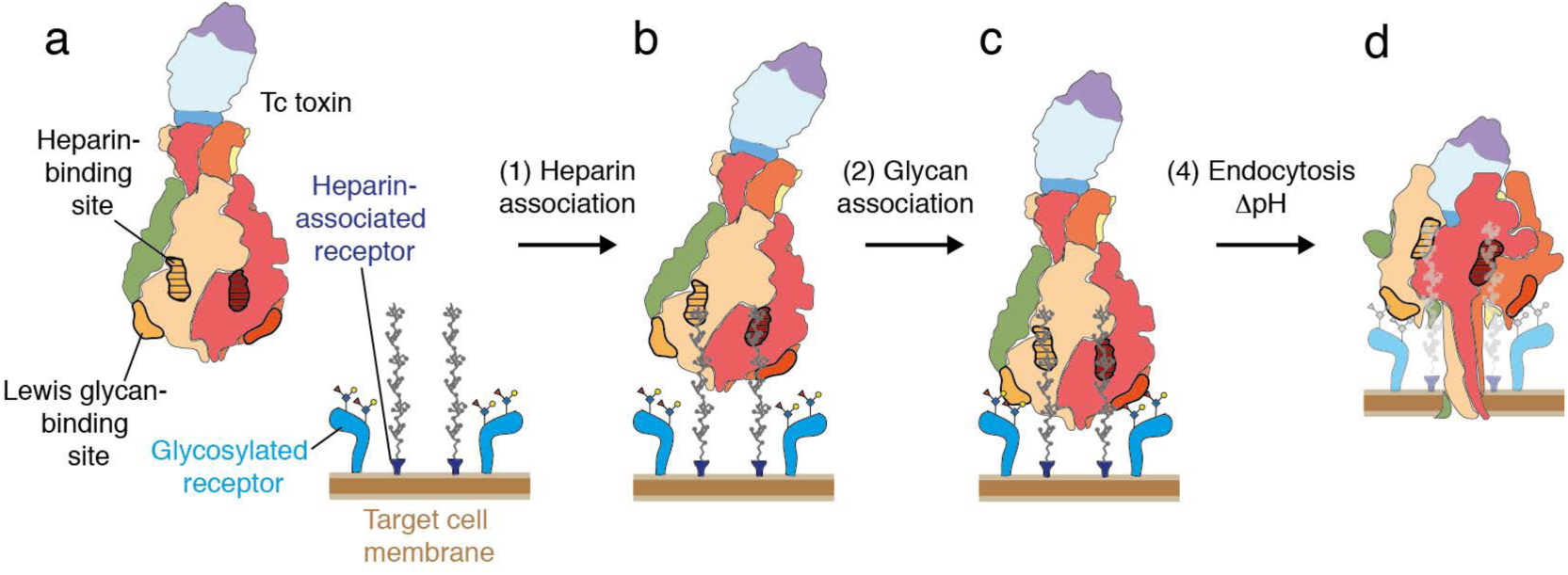
Model of the intoxication mechanism of Tc. a: Tc toxin and target cells with multiple glycosylated and heparin-associated receptors. The binding sites for heparin and Lewis antigens are indicted on two protomers. b: Tc toxin associates to heparin-associated receptors on target cells. c: In a second association reaction, Tc toxin binds to additional glycosylated receptors, resulting in a tighter toxin-cell-complex. d: Upon endocytosis, the shift of pH to acidic values induces prepore-to-pore transition and membrane permeation of the Tc toxin.

## Acknowledgements

We thank Dr. O. Hofnagel and Dr. D. Prumbaum for excellent assistance in electron microscopy and acquisition of the datasets. We thank A. Elsner and K. Vogel-Bachmayr for purification of Pl-TcdA1, Mm-TcdA4, and Pl-TcdB2-TccC3, respectively, and M. Hülseweh for preparation of HEK cells. The authors would like to thank Dr. T. Wagner for aiding with Fiji, and M. Schulz for helping with flow cytometry. This work was supported by funds from the Max Planck Society (to S.R. and P.H.S.) and the European Research Council under the European Union’s Seventh Framework Programme (FP7/2007-2013) (grant no. 615984) (to S.R.).

## Author Contributions

S.R. and P.H.S. designed the project. D.R., O.S and F.L. prepared fluorescently labeled TcAs for the glycan array and cellular experiments. F.B and P.K. carried out the glycan arrays and interpreted the data. F.B. prepared the BSA-Lewis X adduct. D.R. analyzed the affinities of proteins to glycans, performed flow cytometry experiments, prepared cryo-EM samples, processed and analyzed the cryo-EM data. O.S. recorded and analyzed confocal microscopy images. D.R., O.S., F.B. and P.K. prepared the figures. All authors discussed the results and contributed in writing of the manuscript.

## Author Information

The coordinates for the EM structures have been deposited in the Electron Microscopy Data Bank under accession number ….. Correspondence and requests for materials should be addressed to S.R.

## Methods

### Protein production

*P. luminescens* TcdB2-TccC3 and Pl-TcdA1 were expressed in BL21-CodonPlus(DE3)-RIPL in 10 L LB medium and purified as described previously^12^. The other TcAs (Xn-XptA1, Mm-TcdA4 and Yp-TcaATcaB were expressed in BL21-CodonPlus(DE3)-RIPL and purified as described previously^23^. The expression plasmid for PNGase F, pOPH6, was a gift from Shaun Lott (Addgene plasmid #40315) and PNGase F was expressed and purified as described previously^39^.

### Labeling of TcAs for the glycan interaction screen

Pl-TcdA1 was labeled with AlexaFluor488-C5-maleimide (Thermo Fisher, Cat. No. A10254) in a 1:3 molar ratio in 20 mM HEPES-NaOH pH 7.3, 150 mM NaCl, 0.05% Tween20 overnight at 4 °C, resulting in fl-TcdA1. The other TcAs (Mm-TcdA4, Xn-XptA1, Yp-TcaATcaB) were labeled with AlexaFluor488 NHS ester (Thermo Fisher, Cat. No. A20000) in a 1:3 molar ratio in 20 mM HEPES-NaOH pH 7.3, 150 mM NaCl, 0.05% Tween-20 overnight at 4 °C. Subsequently, unreacted labelling dye was removed via size exclusion chromatography (SEC) on a Superose 6 Increase column (GE Healthcare Life Sciences). All labeled proteins were flash-frozen in liquid nitrogen in the presence of 35% glycerol and stored at −80 °C until usage.

### Glycan interaction microarray

Amine-functionalized oligosaccharides or natural heparin (5 kDa) isolated from porcine intestinal mucosae (Sigma-Aldrich) were immobilized on commercial *N*-hydroxysuccinimide (NHS) ester-activated microarray slides (CodeLink Activated Slides; SurModics, Eden Prairie, MN, USA) using a piezoelectric spotting device (S3; Scienion, Berlin, Germany). Initial screening was performed with a comprehensive microarray displaying a large library of oligosaccharide antigens (Supplementary Table 2), as described before^40^. The HS/heparin microarray has been published before^41, 42^ (Supplementary Table 3). Other focused microarray slides that were used to confirm initial hits contained oligosaccharides related to Lewis X. Samples for spotting were diluted in 50 mM sodium phosphate buffer, pH 8.5. Spotted slides were incubated in a humid chamber for 24 h at room temperature to complete coupling reactions. Remaining NHS ester groups were deactivated with 50 mM ethanolamine in 50 mM sodium phosphate buffer, pH 9, for 1 h at 50 °C. Slides were rinsed three times with deionized water, dried by centrifugation (300 x *g*, 5 min) and stored desiccated until use. Spotted and quenched microarray slides were blocked using 100 mM (4-(2-hydroxyethyl)-1-piperazineethanesulfonic acid (HEPES) pH 7.4 with 200 mM NaCl and 1 % (w/v) BSA (PBS-BSA) for 1 h at room temperature, washed three times with 100 mM HEPES pH 7.4 with 200 mM NaCl and 0.05 % (v/v) Tween-20 (wash buffer) and dried by centrifugation. FlexWell 16 or FlexWell 64 grids (Grace Bio-Labs, Bend, OR, USA) were applied to the slides to yield 16 or 64 wells for individual experiments, depending on the printing patterns. Slides were incubated with samples diluted in 100 mM HEPES pH 7.4 with 200 mM NaCl, 0.01 % (v/v) Tween-20 and 1 % (w/v) BSA for 1 h at room temperature in a humid chamber. Wells were washed three times using wash buffer and rinsed once with deionized water. Grids were removed and slides were dried by centrifugation (300 x *g*, 5 min). The microarray slides were scanned with a GenePix 4300A scanner (Molecular Devices; Sunnyvale, CA, USA). The photomultiplier tube voltage was adjusted to reveal scans free of saturated signals. Image analysis was carried out with the GenePix Pro 7 software supplied with the instrument. Background-subtracted mean fluorescence intensity (MFI) values were exported for further analysis.

### Conjugation of Lewis X and BSA

We prepared a glycoconjugate composed of Lewis X trisaccharide (compound **7**, Figure 2c) and BSA following modified procedures described before^43^. The procedure is illustrated in Supplementary Figure 2b. First, **7** was combined with a 6-fold molar excess of di-*p*-nitrophenyl adipate in 400 µL anhydrous dimethyl sulfoxide (DMSO)/pyridine (2:1) and 10 µL triethylamine (Et_3_N). The reaction mixture was incubated for 2 hours at room temperature while stirring. Solvents were evaporated by lyophilization. Dried reaction products were successively washed with dichloromethane and chloroform (ten times 1 mL each) until thin-layer chromatography (TLC) revealed complete removal of non-reacted crosslinker. The washed reaction products were solubilized in DMSO, transferred to new reaction tubes and lyophilized. Dried products were reacted with bovine serum albumin (BSA; Pan Biotech) in 100 mM sodium phosphate buffer, pH 8, for 24 h at room temperature while stirring. The resulting glycoconjugate was washed and concentrated with deionized water using 10 kDa centrifugal filter units (Merck Millipore, Tullagreen, Ireland).

Conjugation was confirmed by MALDI-TOF MS using an Autoflex Speed system (Bruker Daltonics; Bremen, Germany). Samples were spotted using the dried droplet technique with 2,5-dihydroxyacetophenone (DHAP) as matrix on MTP 384 ground steel target plates (Bruker Daltonics). Samples were prepared by mixing 2 µl of desalted protein sample with 2 µL of DHAP matrix and 2 µL of 2 % (v/v) trifluoroacetic acid (TFA) prior to spotting. The mass spectrometer was operated in linear positive mode. Mass spectra were acquired over an m/z range from 30,000 to 210,000 and data was analyzed with the FlexAnalysis software provided with the instrument. The mass shift of about 14,500 Da showed that, on average, 20 molecules of Lewis X were conjugated per molecule of BSA (Supplementary Figure 2c).

### Biolayer interferometry (BLI)

The affinity of Pl-TcdA1 to BSA-Lewis X was determined by BLI using an OctedRed 384 (forteBio, Pall Life Sciences) and streptavidin biosensors. BSA-Lewis X was biotinylated in 20 mM HEPES-NaOH pH 7.3, 150 mM NaCl, 0.05% Tween20 (labelling buffer) with NHS-LC-Biotin (Thermo Fisher, Cat. No. 21336) in a 1:3 molar ratio for 2 h at room temperature. Unreacted biotin label was removed via SEC on a Superdex 200 Increase column (GE Healthcare Life Sciences). Biotinylated BSA-Lewis X was immobilized on streptavidin biosensors at 10 µg/mL, and Pl-TcdA1 pentamer concentration in solution was 25 – 1600 nM. BLI sensorgrams were measured in three steps: baseline (300 s), association (90 s), and dissociation (60 s). The sensorgrams were corrected for background association of Pl-TcdA1 on unloaded streptavidin biosensors. On-and off-rates of TcA binding were determined simultaneously by a global curve fit according to a 1:1 binding model. All BLI steps were performed in labelling buffer with 0.3 mg/mL BSA.

The affinity of Mm-TcdA4 to heparin was determined analogously using the same instrument setup and buffers as described above. Biotinylated porcine intestinal mucosa heparin (Merck Millipore) was immobilized on streptavidin biosensors at 10 µg/mL, and Mm-TcdA4 pentamer concentration in solution was 8 – 500 nM. BLI sensorgrams were measured with 600 s association and 600 s dissociation time, and corrected for background association of Mm-TcdA4 on unloaded streptavidin biosensors. On-and off-rates of TcA binding were determined simultaneously by a global curve fit according to a 1:1 binding model.

### Deglycosylation of HEK293T cells

5.5 *×* 10^6^ HEK 293T cells (Thermo Fisher) attached to 10 cm cell culture dishes (Sarstedt) in DMEM/F12 + 10% FBS medium were incubated with 20 μg PNGase F (self-prepared for flow cytometry, NEB Cat. No. P0704 for confocal microscopy) for 5 h at 37 °C. Subsequently, cells were detached from the surface by carefully resuspending them, washed once in DMEM/F12 + FBS and resuspended in 1.8 mL PBS immediately before subsequent intoxication and flow cytometry.

### Flow cytometry

Immediately after deglycosylation and washing, HEK293T or HEK293 GnTI^-^ cells (Thermo Fisher) were incubated with 200 nM (monomer concentration) TcdA1 labeled with AlexaFluor488 for 15 min. Subsequently, cells were washed one time, resuspended in PBS without toxin and injected into a LSRII flow cytometer (BD Biosciences). 50000 cells were counted in total. Cell fluorescence caused by TcdA1 binding was analyzed using FlowJo.

### Intoxication assay

HEK 293T cells or HEK 293 GnTI^-^ cells were intoxicated with pre-formed holotoxin formed by Pl-TcdA1 and PL-TcdB2–TccC3. Cells (5 × 10^4^) were grown adherently in 400 µL DMEM/F12 medium (Pan Biotech) overnight and 0.5 or 2 nM holotoxin was subsequently added. Incubation was performed for 16 h at 37 °C before imaging. Experiments were performed in triplicate. Cells were not tested for *Mycoplasma* contamination.

### Confocal microscopy

Prior to confocal microscopy experiments, cells (HEK 293T and HEK 293 GnTI^-^) were seeded at a density of 0.3 *×* 10^6^ cells per dish and grown on 35 mm poly-d-lysine coated glass-bottom culture dishes (MatTek) until 30-40% confluency in 2 mL DMEM/F12 + 10 % FBS for 36 hours in a 37 °C, 5 % CO_2_ humidified atmosphere. Confocal imaging was carried out at 37 °C in a 5 % CO_2_ atmosphere using a ZEISS LSM 800 microscope equipped with a C-Apochromat 40x/1.2 W objective. After the initial image was taken (0 h), AlexaFluor488 labelled TcdA1 was added to a final concentration of 500 pM. Images were then taken at 10-minute intervals for 8 hours. The experiments were done in three biological replicates for each cell type.

Images taken at 1 hour intervals were processed in Fiji^44^. A cell mask was generated by using the Trainable Weka Segmentation plugin^45^ on the transmitted light channel and then used to focus on the relevant areas of the fluorescent channel. The mean grey values from the fluorescent channel were then measured for each image, which represent the sum of all grey pixel values (fluorescence) in the selections divided by the total number of pixels (cell surface). The autofluorescence for each replicate was removed by subtracting the mean grey value of the initial 0 h image from later post-intoxication images. The averages and standard deviation for this normalized mean fluorescence intensity were then calculated and plotted.

### Crosslinking of TcdA1 and BSA-Lewis X

TcdA1 (1 μM) and BSA-Lewis X (30 μM) were incubated in 20 mM HEPES-NaOH pH 8.0, 150mM NaCl, 0.02% Tween-20 overnight at 4 °C. Subsequently, glutaraldehyde was added to a final concentration of 0.1% and proteins were crosslinked for 90 min at 22 °C. The reaction was stopped with 1.7 mM Tris-HCl pH 8.0 and crosslinked TcdA1-BSA-Lewis X was purified from excess of BSA-Lewis X by SEC using a Superose 6 3.2-300 column (GE Healthcare Life Sciences).

### Cryo-EM sample preparation and data acquisition of TcdA1-BSA-Lewis X

4 µL of 0.06 mg/mL crosslinked complex were applied to glow-discharged holey carbon grids (Quantifoil, QF 1.2/1.3, 300 mesh) covered with a 2 nm carbon layer. After 10 min of incubation time on the grid, the grid was manually blotted and another 4 µL were applied and incubated for 20 s. Subsequently, the sample was vitrified in liquid ethane with a Cryoplunge3 (Cp3, Gatan) using 1.8 s blotting time at 90 % humidity.

A dataset of TcdA1-BSA-Lewis X was collected using a Cs corrected Titan Krios equipped with an XFEG and a Falcon III direct electron detector. Images were recorded using the automated acquisition program EPU (FEI) at a magnification of 59,000x, corresponding to a pixel size of 1.11 Å/pixel on the specimen level. 4,992 movie-mode images were acquired in a defocus range of 1.0 to 2.5 µm. Each movie comprised 40 frames acquired over 1.5 s with a total cumulative dose of ∼ 100 e^-^/Å^2^.

### Image processing of TcdA1-BSA-Lewis X

After initial screening of all micrographs, 3878 integrated images were selected for further processing. Movie frames were aligned, dose-corrected and averaged using MotionCor2^46^. CTF parameters were estimated using CTER^47^, implemented in the SPHIRE software package^30^. Initially, 4300 particles were manually picked and 2D class averages used as an autopicking template were generated using ISAC^48^ in SPHIRE. 711,872 particles were auto-picked from the images using Gautomatch (http://www.mrc-lmb.cam.ac.uk/kzhang).

Reference-free 2D classification and cleaning of the dataset was performed with the iterative stable alignment and clustering approach ISAC in SPHIRE. ISAC was performed with a pixel size of 5.61 Å/pixel on the particle level. The ‘Beautifier’ tool of SPHIRE was then applied to obtain refined and sharpened 2D class averages at the original pixel size, showing toxin particles with visible additional density at the TcA pentamer, which corresponds to BSA-Lewis X (Figure 3a). 199,038 particles were selected for subsequent 3D refinement. We applied our previous reconstruction of *P. luminescens* TcdA1 (EMD-3645) as initial model after scaling and filtering it to 25 Å resolution and performed an initial 3D refinement in SPHIRE using Meridien with C5 symmetry. The density corresponding to the bound BSA-Lewis X appears only at very low binarization threshold (Supplementary Figure 4). Therefore, we expanded the symmetry of the input particle to C5 via the “symmetrize” option of SPHIRE, resulting in 995,190 particles. Subsequently, we applied 3D sorting (Sort3D in SPHIRE) to the symmetrized stack using a focused mask that comprises the lower part of the shell and the additional densities. The resulting 3D classes showed 1 – 2 additional densities corresponding to BSA-Lewis X at different orientations. We next rotated the projection parameters of the classes by +72° or −72° in order to orient the most pronounced additional density to the same direction in all classes (Supplementary Figure 4). Finally, after import of the rotated projection parameters and stack de-symmetrization, we ran a local refinement in Meridien with C1 symmetry. The resolution of the final density was estimated to 6.99 / 5.01 Å according to FSC 0.5 / 0.143 after applying a soft Gaussian mask. The B-factor was set to −80.0 Å^2^ for sharpening.

### Cryo-EM sample preparation and data acquisition of TcdA4 and heparin

Incubation of Mm-TcdA4 with porcine mucosa heparin in solution resulted in precipitation. Therefore, 3 µL of 0.10 mg/mL TcdA4 in 20 mM Tris-HCl pH 8.0, 250 mM NaCl, 0.05% Tween-20 was pre-applied on a glow-discharged holey carbon grid (Quantifoil, QF 2/1, 300 mesh) covered with a 2 nm carbon layer. After 20 s of incubation time on the grid, the grid was manually blotted and 3 µL of 0.3 mg/mL heparin from porcine intestinal mucosa (Millipore, Cat. No. 375095) applied and incubated for 2 min at 22 °C. Subsequently, the sample was vitrified in liquid ethane with a Cryoplunge3 (Cp3, Gatan) using 1.8 s blotting time at 89 % humidity.

A dataset of TcdA1-LewisX-BSA was collected at the Max Planck Institute of Molecular Physiology, Dortmund using a Cs corrected Titan Krios equipped with an XFEG and a Falcon III direct electron detector. Images were recorded using the automated acquisition program EPU (FEI) at a magnification of 59,000x, corresponding to a pixel size of 1.11 Å/pixel on the specimen level. 5549 movie-mode images were acquired in a defocus range of 1.2 to 2.2 µm. Each movie comprised 40 frames acquired over 1.5 s with a total cumulative dose of ∼ 100 e^-^/Å^2^.

### Image processing of Mm-TcdA4-heparin

After initial screening of all micrographs, 4749 integrated images were selected for further processing. Movie frames were aligned, dose-corrected and averaged using MotionCor2 ^46^. The integrated images were also used to determine the contrast transfer function (CTF) parameters with CTER^47^, implemented in the SPHIRE software package ^30^. Initially, 4,300 particles were manually picked and 2D class averages used as an autopicking template were generated using ISAC ^48^ in SPHIRE. 477,602 particles were auto-picked from the images using Gautomatch (http://www.mrc-lmb.cam.ac.uk/kzhang).

Reference-free 2D classification and cleaning of the dataset were performed with the iterative stable alignment and clustering approach ISAC in SPHIRE. ISAC was executed with a pixel size of 5.61 Å/pixel on the particle level. The ‘Beautifier’ tool of SPHIRE was then applied to obtain refined and sharpened 2D class averages at the original pixel size, showing high-resolution features (Supplementary Figure 6b). From the initial set of particles, the clean set used for 3D refinement contained 182,506 particles. We applied our previous 3.3 Å reconstruction of *M. morganii* TcdA4 ^23^ (EMD-10035) as an initial model after scaling and filtering it to 25 Å resolution and performed 3D refinement in SPHIRE using Meridien with imposed C5 symmetry. The resolution of the final density was estimated to be 3.6 / 3.2 Å according to FSC 0.5 / 0.143 after applying a soft Gaussian mask. The B-factor was estimated to be 127.6 Å^2^. Local FSC calculation was performed using the Local Resolution tool in SPHIRE. (Supplementary Figure 6e) and the electron density map was filtered according to its local resolution using the 3-D Local Filter tool in SPHIRE.

The local resolution of the additional density corresponding to heparin was however not sufficient to obtain a molecular model. We therefore filtered the map obtained here and the *M. morganii* TcdA4 map without heparin to 5 Å resolution, normalized the maps and subtracted the filtered and normalized TcdA4 map without heparin from the map with heparin. The difference density was filtered to 15 Å resolution to visualize the density corresponding to heparin (Supplementary Figure 7a).

### Molecular docking

We used AutoDock Vina^49^ via the UCSF Chimera plugin to visualize potential conformations of heparin bound to Mm-TcdA4 and Lewis X bound to Pl-TcdA1. Before docking, the interface regions of the proteins were protonated using H++^50^ at pH 7.0 and 150 mM ionic strength. For Mm-TcdA4, we docked a heparin pentasaccharide (molecule code NTP) after geometry optimization in phenix.elbow^51^ to the outer shell region of (residues 100 – 310 and 443 – 984). We chose the conformation that fitted best into the additional density from the obtained docking results. The resulting score of the docking pose was −6.2 kcal/mol. For Pl-TcdA1, we docked the Lewis X trisaccharide to RBD D (residues 1638-1754). We chose two representative conformations which are closely located to the additional density with scores of −4.5 kcal/mol each.

## Supplementary Figures

**Supplementary Figure 1:**
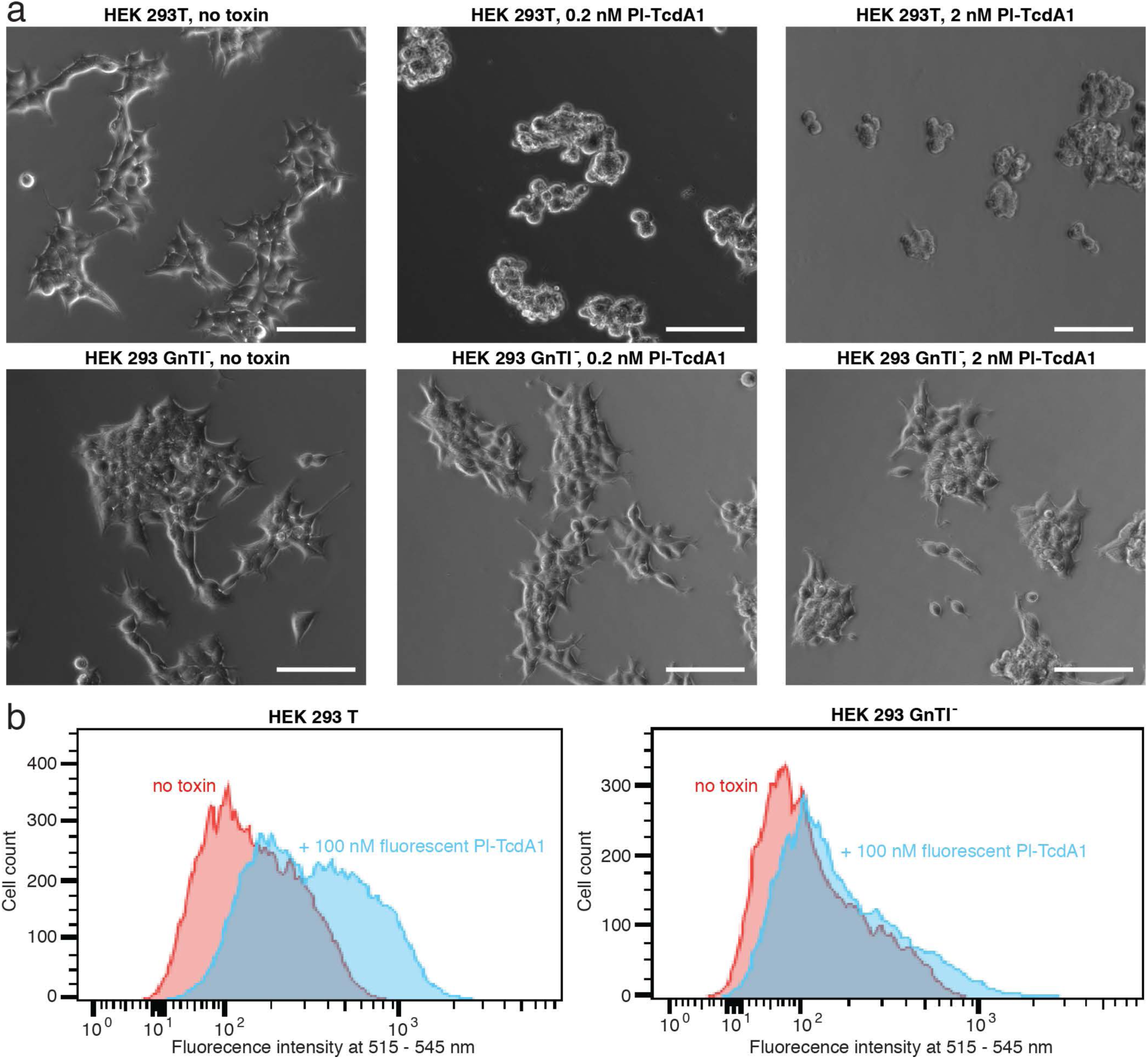
Toxicity and binding of Tc to HEK 293 T and HEK 293 GnTI^-^ cells. a: Intoxication of HEK 293T (top row) and HEK 293 GnTI^-^ (bottom row) with Tc holotoxin (Pl-TcdA1 with Pl-TcdB2-TccC3). Images show cells 16 h after intoxication. Intoxicated cells round up and detach from the surface. Experiments were performed in triplicates with qualitatively identical results. Scale bars, 100 µm. b: Flow cytometry of HEK 293T (left) and HEK 293 GnTI^-^ (right) exposed to AlexaFluor488 labelled Pl-TcdA1. Histograms of cells, both with and without 100 nM Pl-TcdA1 are shown in comparison.

**Supplementary Figure 2:**
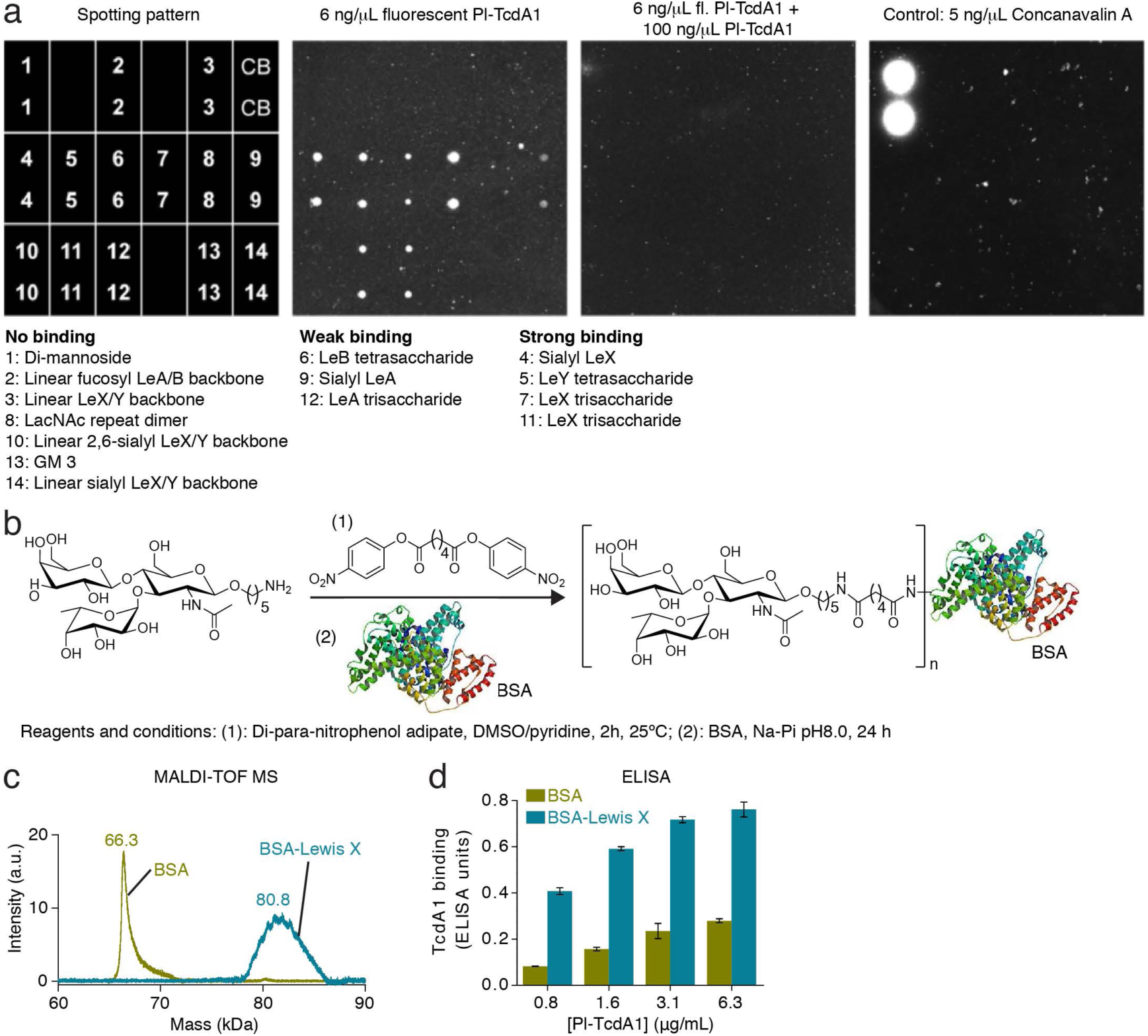
Glycan microarray of Pl-TcdA1 and preparation of BSA-Lewis X. a: Spotting pattern of glycans on the chip surface in duplicates (left), fluorescence readout after incubation with 6 ng/µL fluorescently labeled Pl-TcdA1 (fl-TcdA1) alone (middle left) and together with a 17-fold excess of unlabeled Pl-TcdA1 (middle right), and fluorescence readout after incubation with concanavalin A (right). The glycans immobilized on the chip (structures are shown in Figure 2c) are shown below the chip scheme and are grouped according to their interaction with Pl-TcdA1. b: Reaction scheme showing the preparation of BSA-Lewis X. c: Mass spectrometry (MALDI-TOF MS) of the obtained BSA-Lewis X glycoconjugate in comparison to BSA. The average mass increase of 14.5 kDa shows the immobilization of ∼20 Lewis X trisaccharides per BSA molecule. d: ELISA of non-fluorescent Pl-TcdA1 (0.8 – 6.3 μg/mL) with immobilized BSA-Lewis X or BSA, showing dose-dependent binding of Pl-TcdA1 to BSA-Lewis X and weak binding to BSA.

**Supplementary Figure 3:**
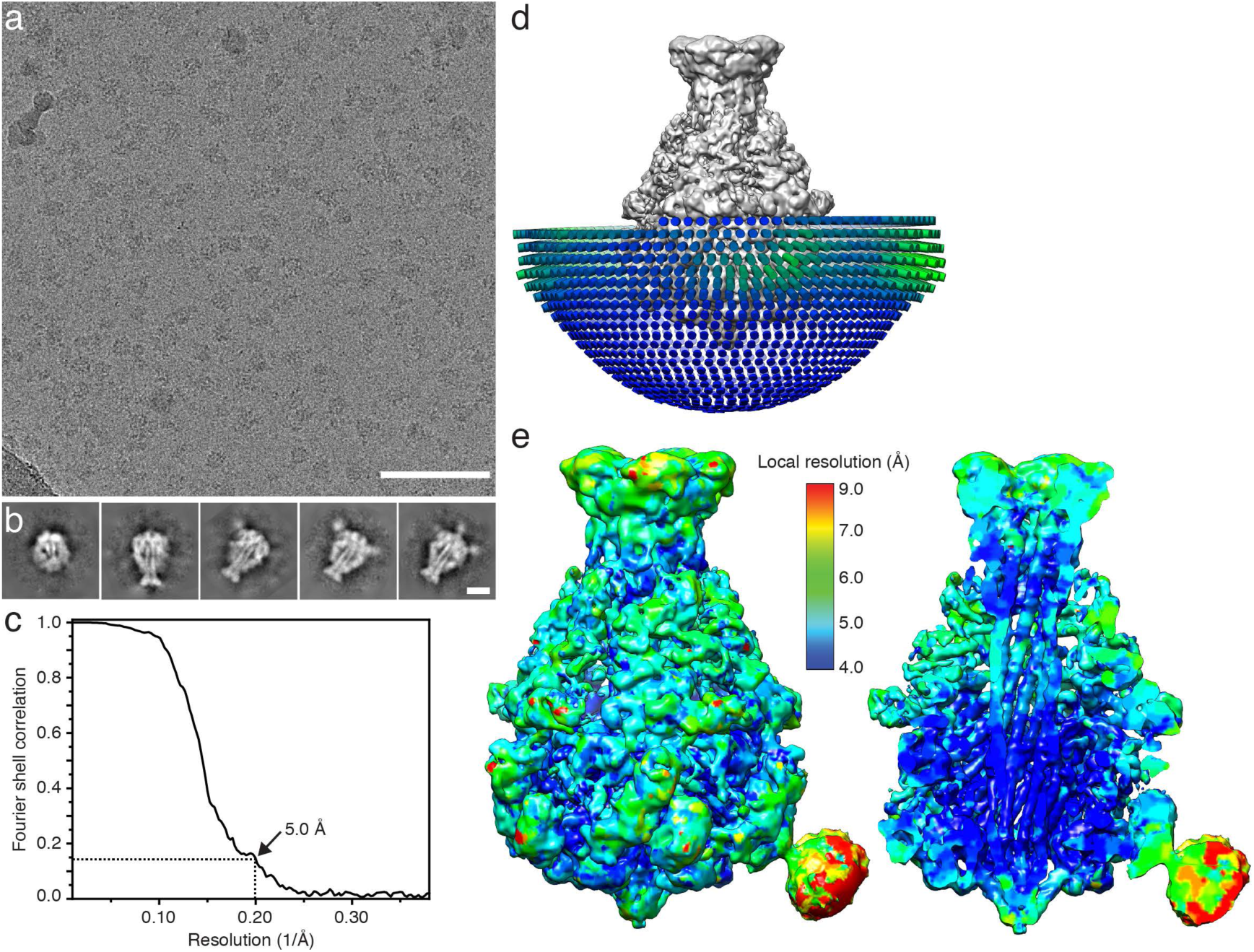
Cryo-EM of Pl-TcdA41 and crosslinked BSA-Lewis X. a: Typical digital micrograph area of vitrified Pl-TcdA1-BSA-Lewis X complexes at a defocus of 2 μm and a total dose of 100 e^−^ Å^−2^ acquired with a Falcon III direct electron detector. Scale bar, 100 nm. b: Representative reference-free 2D class averages obtained by ISAC and subsequently resampled to the original pixel size, refined and sharpened, using the Beautifier tool implemented in the SPHIRE software package. Scale bar, 10 nm. c: Fourier shell correlation (FSC) of the cryo-EM map (black curve). The 0.143 FSC cut-off criterion indicates that the cryo-EM map has an average resolution of 5.0 Å. d: Angular distribution for the final round of the refinement. Each stick represents a projection view. Size and color of the stick is proportional to the number of particles. e: Surface and cross-section of the cryo-EM density map colored according to the local resolution. The map sections corresponding to the Pl-TcdA1 pentamer and BSA-Lewis X are shown at different binarization thresholds.

**Supplementary Figure 4:**
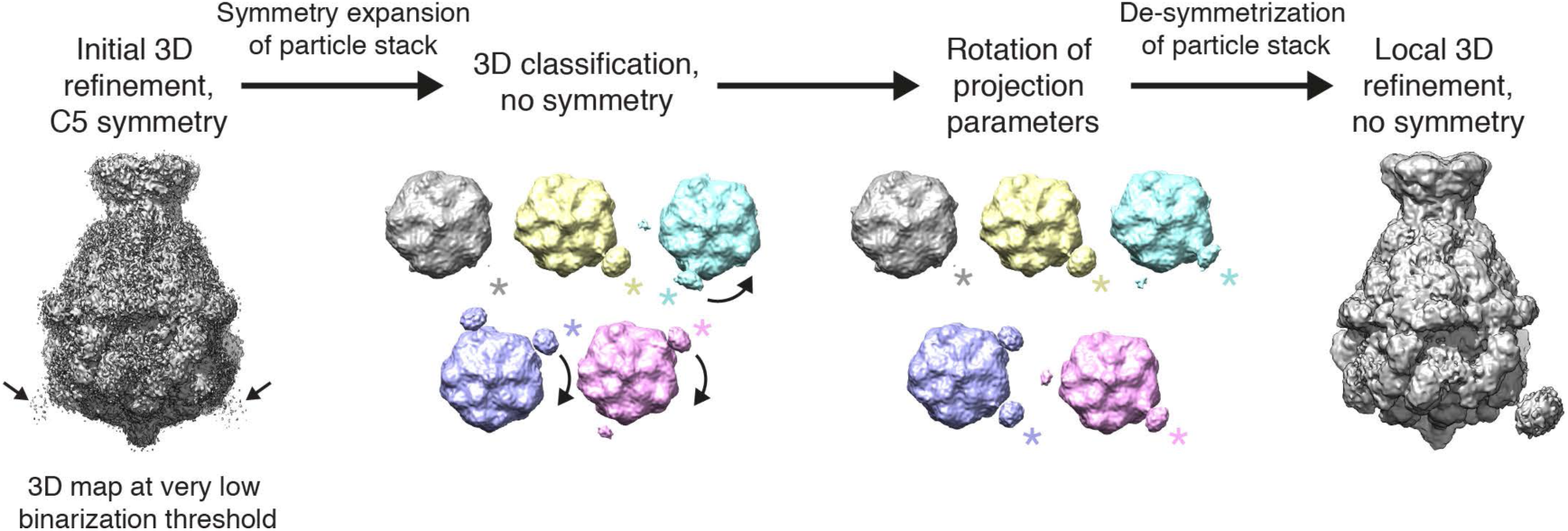
Workflow of 3D classification and refinement for Pl-TcdA1-BSA-Lewis X. Initial 3D refinement with C5 symmetry resulted in weak density for BSA-Lewis X (arrows) which binds sub-stoichiometrically (left panel). Therefore, we performed symmetry expansion of the particle stack and 3D classification after 3D refinement. The 3D classes were rotated in 72° steps so that the additional density corresponding to BSA-Lewis X was located in the same position and the projection parameters adjusted. The stack was then de-symmetrized and a local 3D refinement without symmetry was performed, resulting in a map with BSA-Lewis X oriented at one interaction site of Pl-TcdA1 (right panel). The resolution of the final map is 5 Å.

**Supplementary Figure 5:**
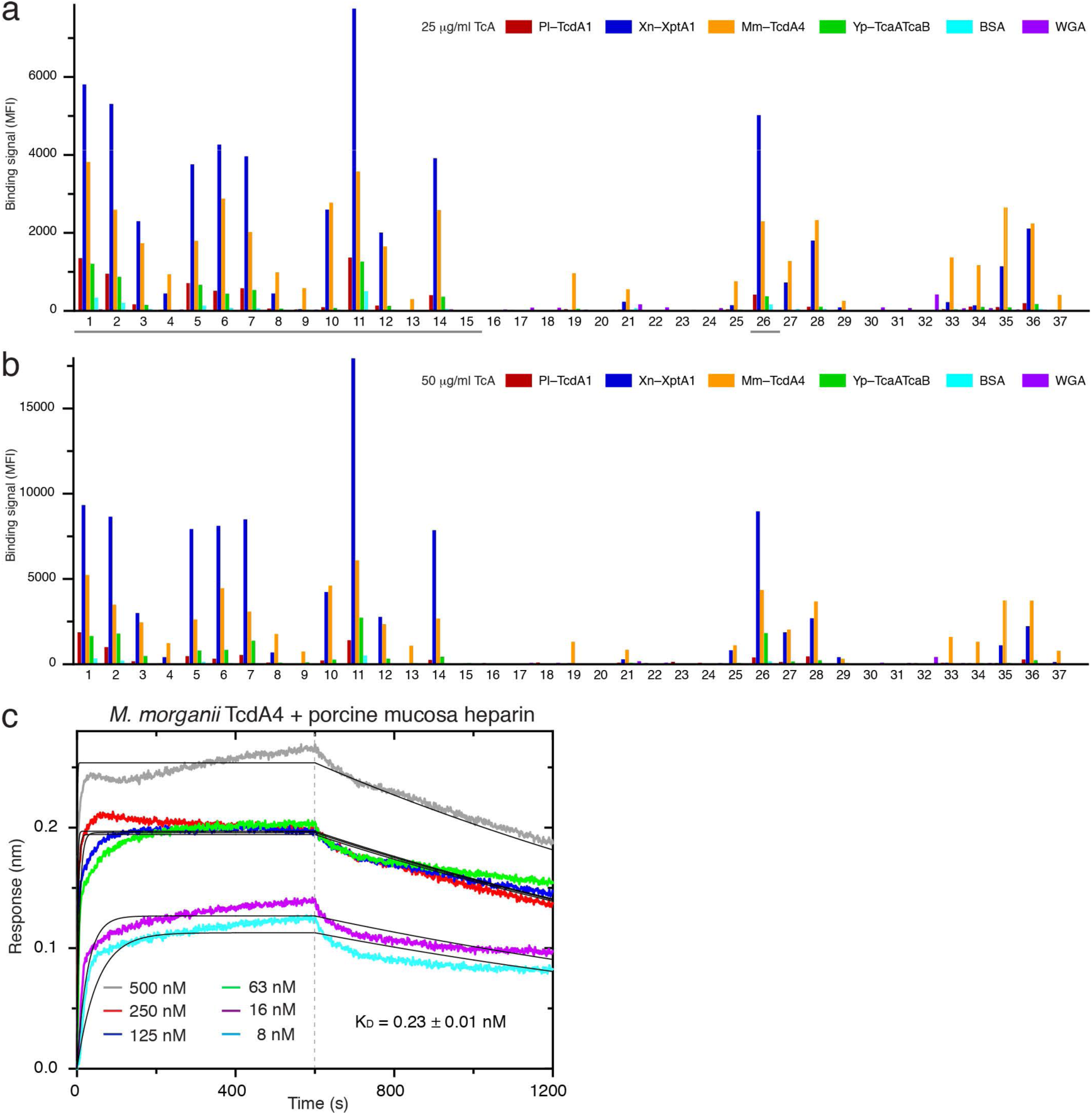
Interaction of TcA with heparins. a,b: Glycan microarray showing the interaction of Pl-TcdA1, Xn-XptA1, Mm-TcdA4, Yp-TcaA-TcaB with various heparins and heparin-like glycans^41, 42^. The protein concentration is 25 μg/ml (a) and 50 μg/ml (b), respectively. BSA and WGA were used as controls. At positions 15 and 37, PBS was spotted on the array as negative control. The gray bars in (a) indicate the molecules that are presented in Figure 4a,b. c: BLI sensorgrams of Mm-TcdA4 (8 nM – 500 nM) with immobilized biotinylated porcine intestinal mucosa heparin. The black lines show a global fit according to a 1:1 binding model, resulting in an apparent K_D_ of 228 ± 3 pM, k_on_ of 2.45 ± 0.03*×*10^4^ M^-1^ s^-1^, and k_off_ of 4.49 ± 0.03*×*10^-4^ s^-1^.

**Supplementary Figure 6:**
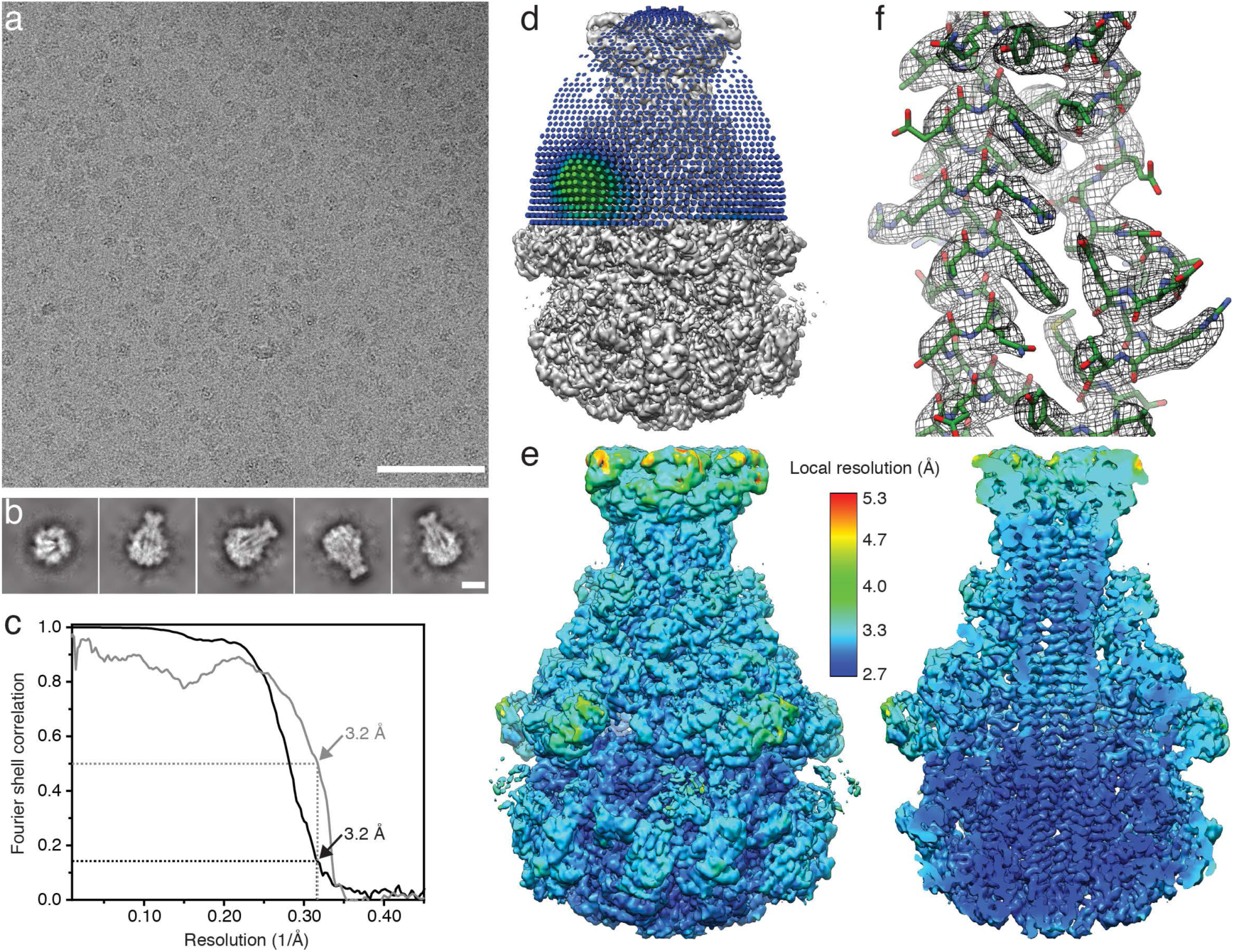
Cryo-EM of Mm-TcdA4 in complex with heparin. a: Typical digital micrograph area of vitrified TcdA4-heparin complexes at a defocus of 2 μm and a total dose of 100 e^−^ Å^−2^ acquired with a Falcon III direct electron detector. Scale bar, 100 nm. b: Representative reference-free 2D class averages obtained by ISAC and subsequently resampled to the original pixel size, refined and sharpened, using the Beautifier tool implemented in the SPHIRE software package. Scale bar, 10 nm. c: Fourier shell correlation (FSC) of the cryo-EM map (black curve). The 0.143 FSC cut-off criterion indicates that the cryo-EM map has an average resolution of 3.2 Å. The gray curve shows the FSC curve between the final map versus the atomic model. The 0.5 FSC cut-off criterion indicates a resolution of 3.2 Å. d: Angular distribution for the final round of the refinement. Each stick represents a projection view. Size and color of the stick is proportional to the number of particles. e: Surface and cross-section of the cryo-EM density map colored according to the local resolution. f: Superimposition of the cryo-EM density map and the model, shown for a representative area in the *α*-helical channel.

**Supplementary Figure 7:**
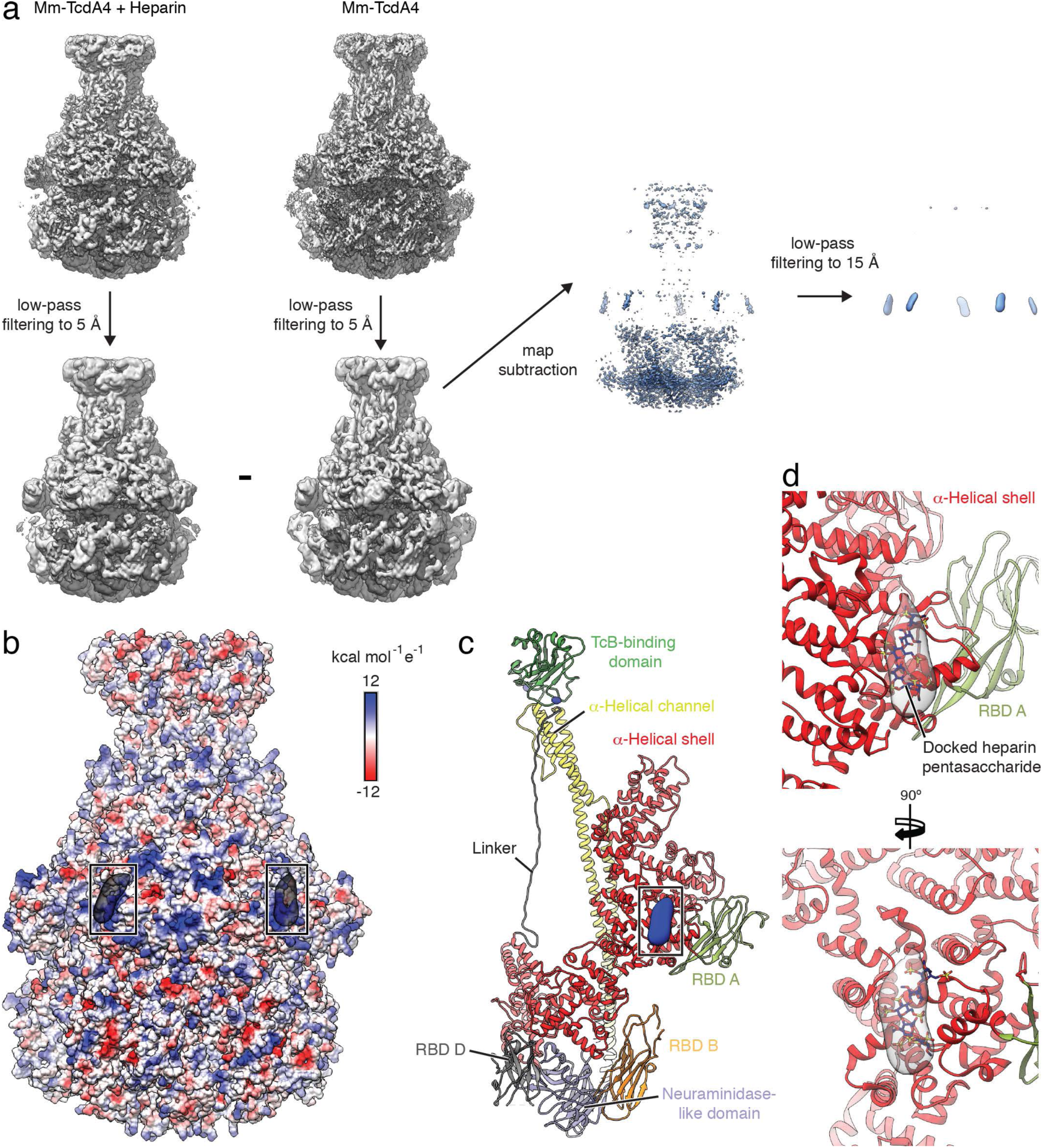
Comparison of Mm-TcdA4 without and with heparin. a: Illustration of map subtraction to obtain the difference density map between Mm-TcdA4 in the absence of heparin ^23^ and Mm-TcdA4 in the presence of porcine intestinal mucosa heparin. After lowpass filtering of both maps to 5 Å, the map of Mm-TcdA4 is subtracted from the map of Mm-TcdA4 with heparin. The obtained difference density map is filtered to 15 Å resolution for illustrative purposes. b: Surface representation of Mm-TcdA4, colored according to the Coulomb potential (kcal mol^-1^ e^-1^) at pH 7.0. The boxes indicate the obtained difference density map (transparent gray) on two protomers. c: Model of one Mm-TcdA4 protomer with difference density map (blue) obtained in panel a. d: Illustration of the docked pentasaccharide (NTP, dark blue) on Mm-TcdA4. The docking solution that resulted in the best match with the difference density (transparent gray) is shown.

## Supplementary Movie legends

Supplementary Movie 1: Cryo-EM density map of Pl-TcdA1/BSA-Lewis X. The map section corresponding to BSA-Lewis X is displayed at a lower binarization threshold. A surface representation of RBD D and the docked BSA-Lewis X molecules are shown.

Supplementary Movie 2: Cryo-EM density map of Mm-TcdA4/heparin with the fitted atomic model. The difference density of the cryo-EM maps of Mm-TcdA4/heparin and Mm-TcdA4 is shown at the heparin binding site at the *α*-helical shell. The docked heparin is shown in the difference density.

## Supplementary Tables

**Supplementary Table 1:** Cryo-EM data collection, refinement and validation statistics for Mm-TcdA4/Heparin and Pl-TcdA1/BSA-Lewis X.

|  | Mm-TcdA4/Heparin | Pl-TcdA1/BSA-Lewis X |
| --- | --- | --- |
| **Data collection and processing** |  |  |
| Camera | Falcon III | Falcon III |
| Magnification | 59,000 | 59,000 |
| Voltage (kV) | 300 | 300 |
| Exposure time (s) | 1.5 | 1.5 |
| Total electron exposure (e-/Å<sup>2</sup>) | 100 | 100 |
| Defocus range (µm) | 1.2 – 2.2 | 1.0 – 2.5 |
| Pixel size (Å) | 1.11 | 1.11 |
| Symmetry imposed | C5 | C5 + C1 |
| Initial particle images (no.) | 477,602 | 711,872 |
| Final particle images (no.) | 182,506 | 199,038 |
| Map resolution (Å) | 3.2 | 5.0 |
| FSC threshold | 0.143 | 0.143 |
| Map sharpening *B* factor (Å<sup>2</sup>) | -127.06 | -80.00 |
| **Refinement** |  |  |
| Initial model used (PDB code) | 6RW9 |  |
| Model resolution (Å) | 3.2 |  |
| FSC threshold | 0.5 |  |
| Model composition |  |  |
| Non-hydrogen atoms | 88,700 |  |
| Protein residues | 11,425 |  |
| Ligands |  |  |
| Mean B factors (Å<sup>2</sup>) |  |  |
| Protein | 55.78 |  |
| R.m.s. deviations |  |  |
| Bond lengths (Å) | 0.011 |  |
| Bond angles (°) | 0.901 |  |
| Validation |  |  |
| MolProbity score | 2.46 |  |
| EMRinger score | 2.22 |  |
| Clashscore | 13.89 |  |
| Rotamer outliers (%) | 2.83 |  |
| Ramachandran plot |  |  |
| Favored (%) | 92.42 |  |
| Allowed (%) | 7.40 |  |
| Disallowed (%) | 0.18 |  |

**Supplementary Table 2:**
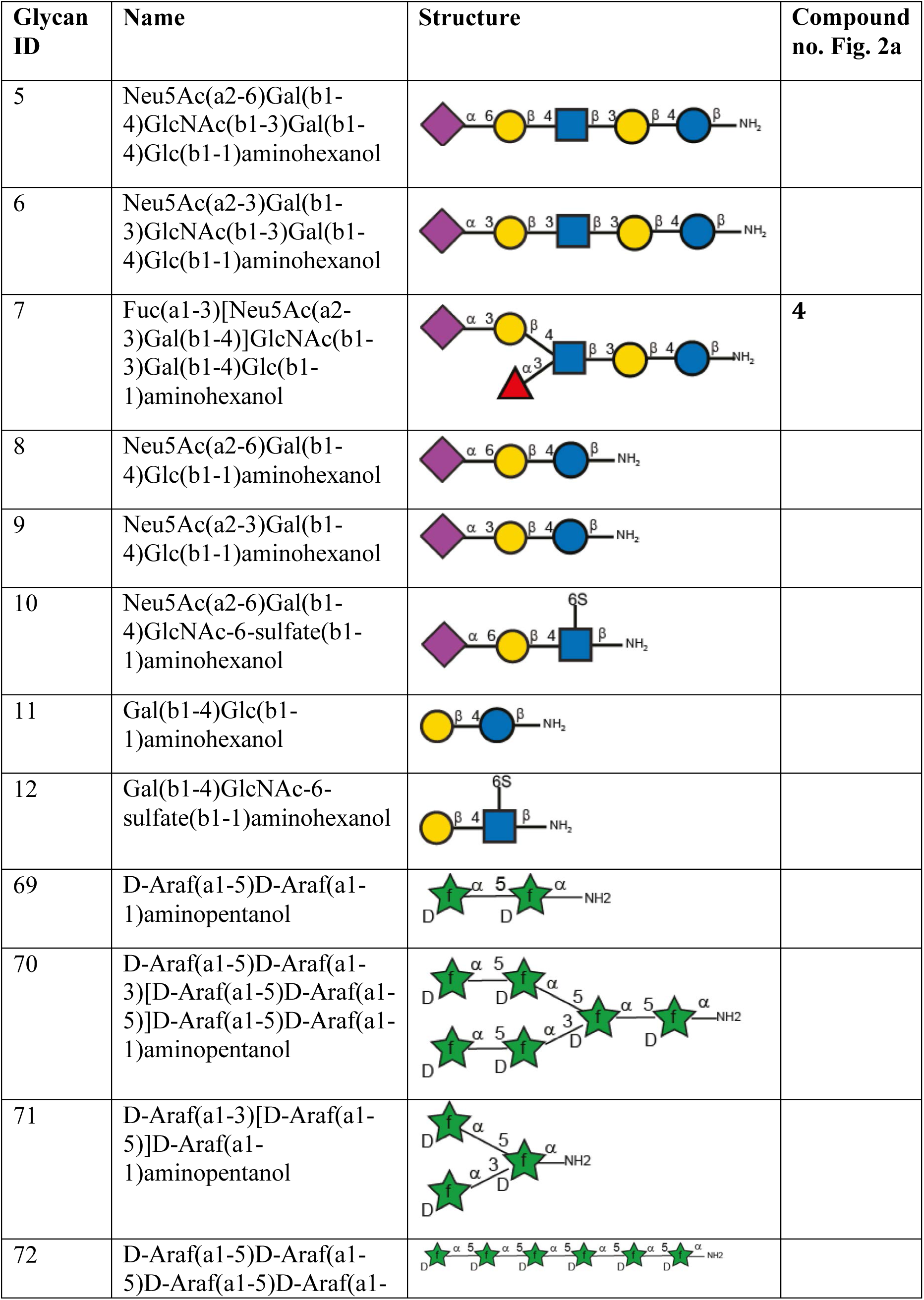
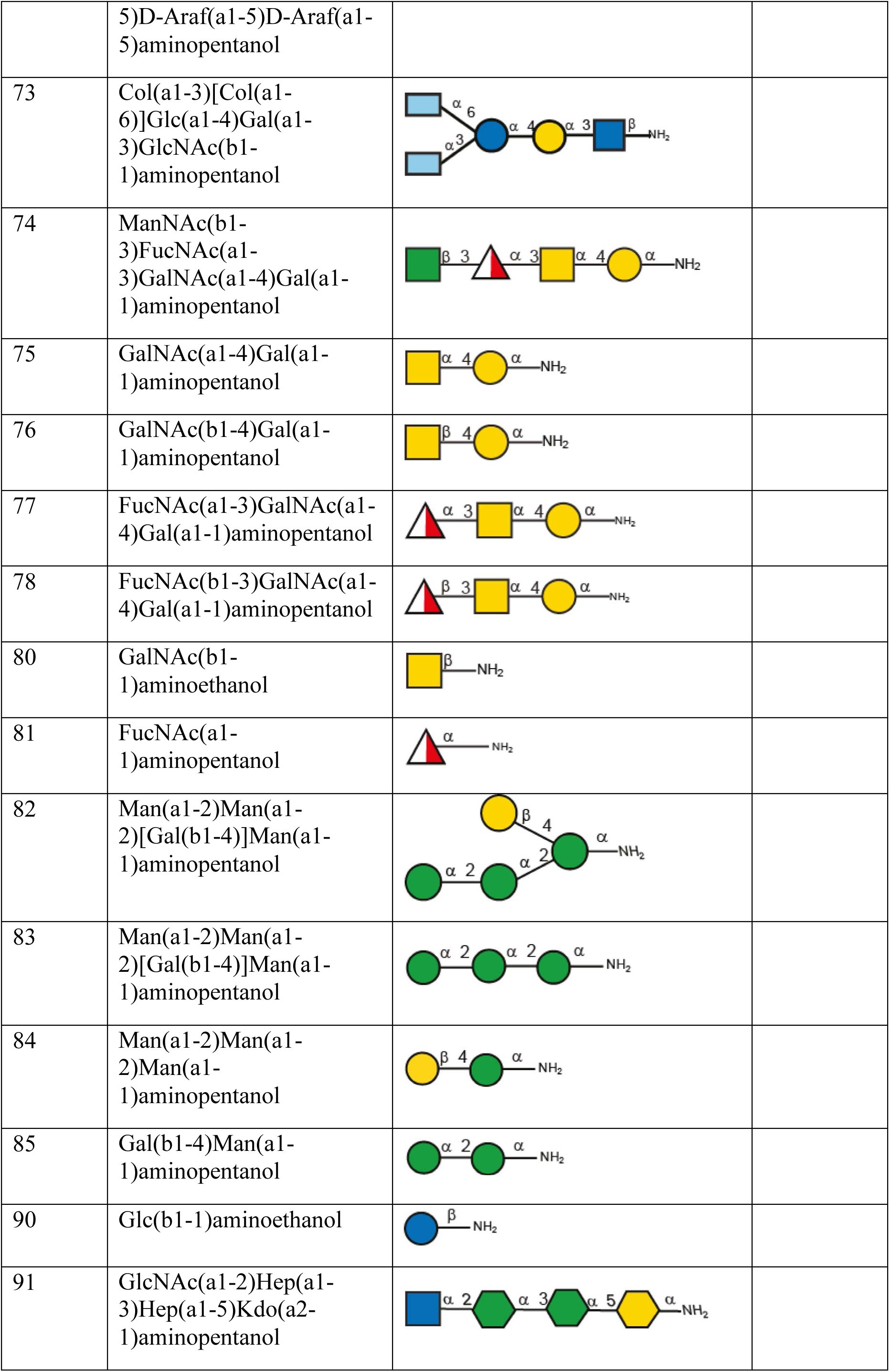
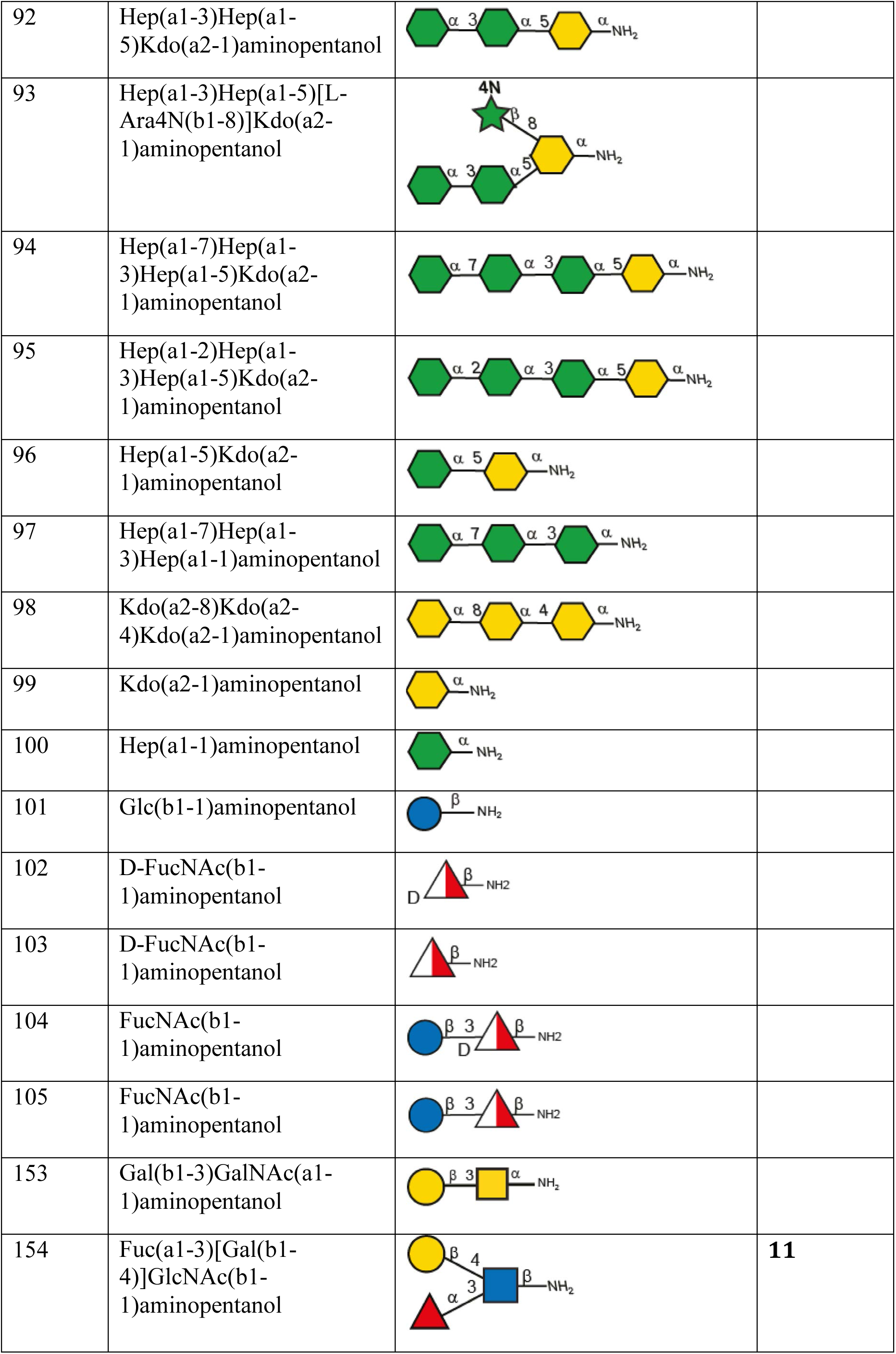
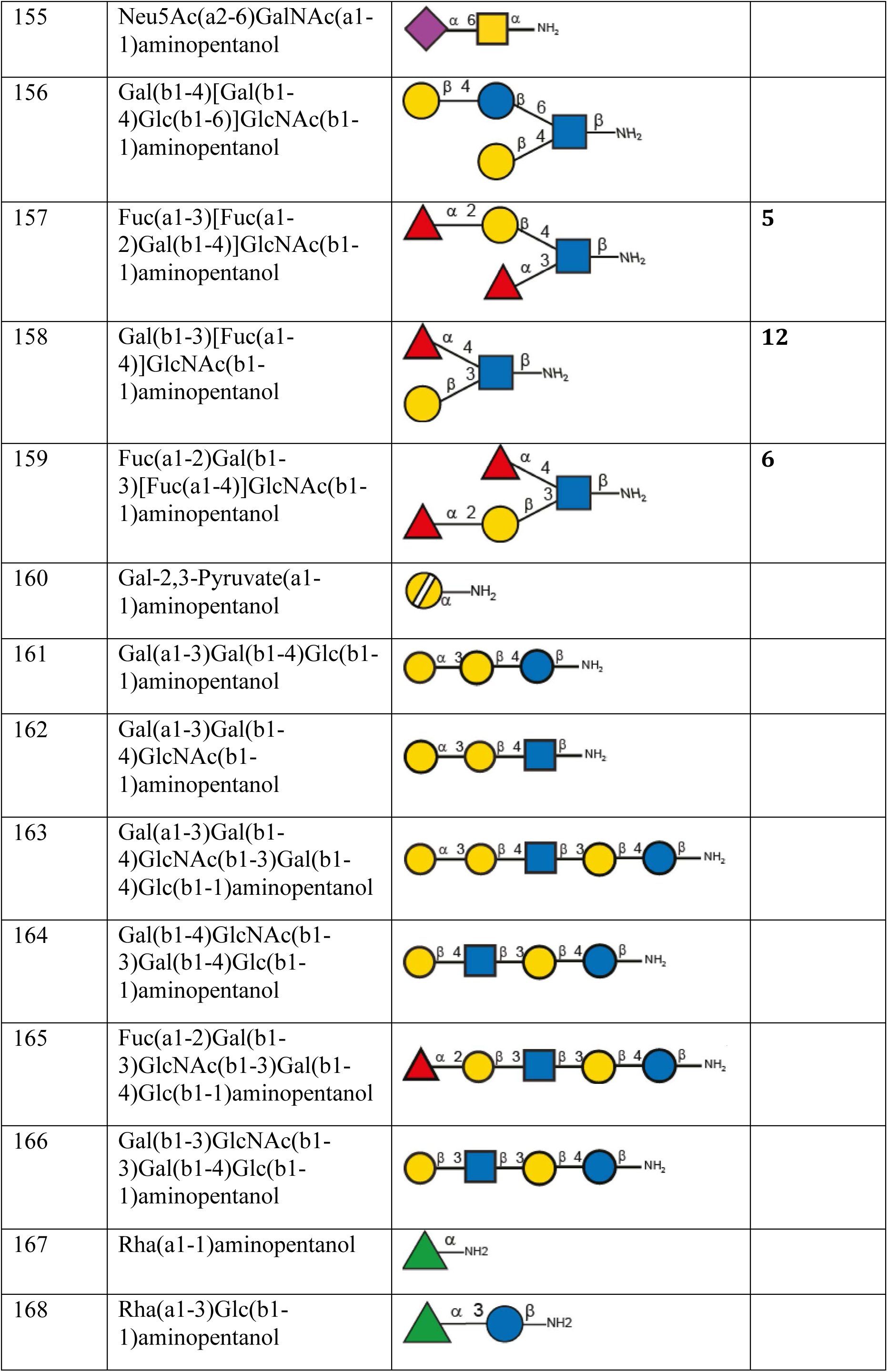
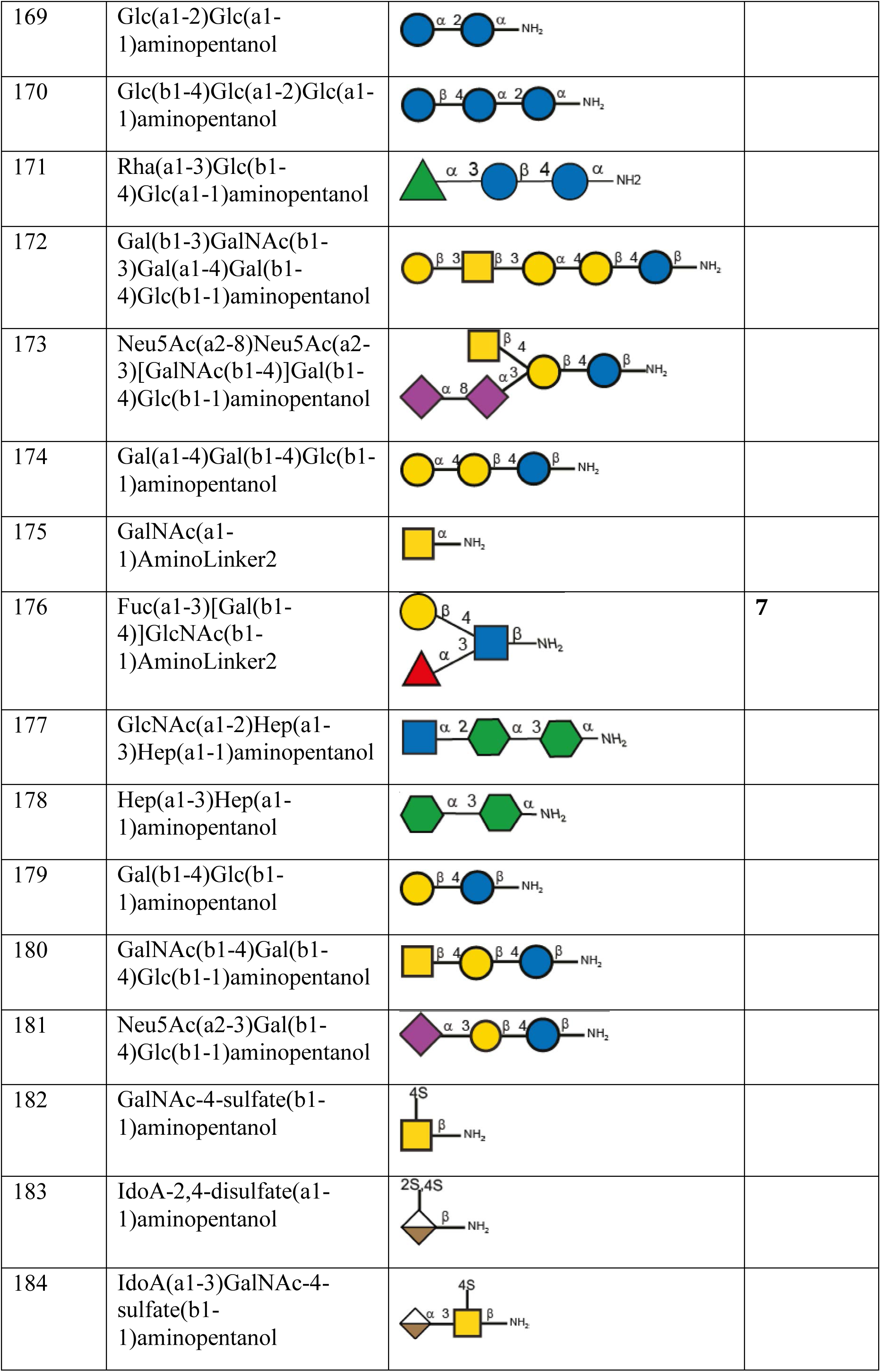
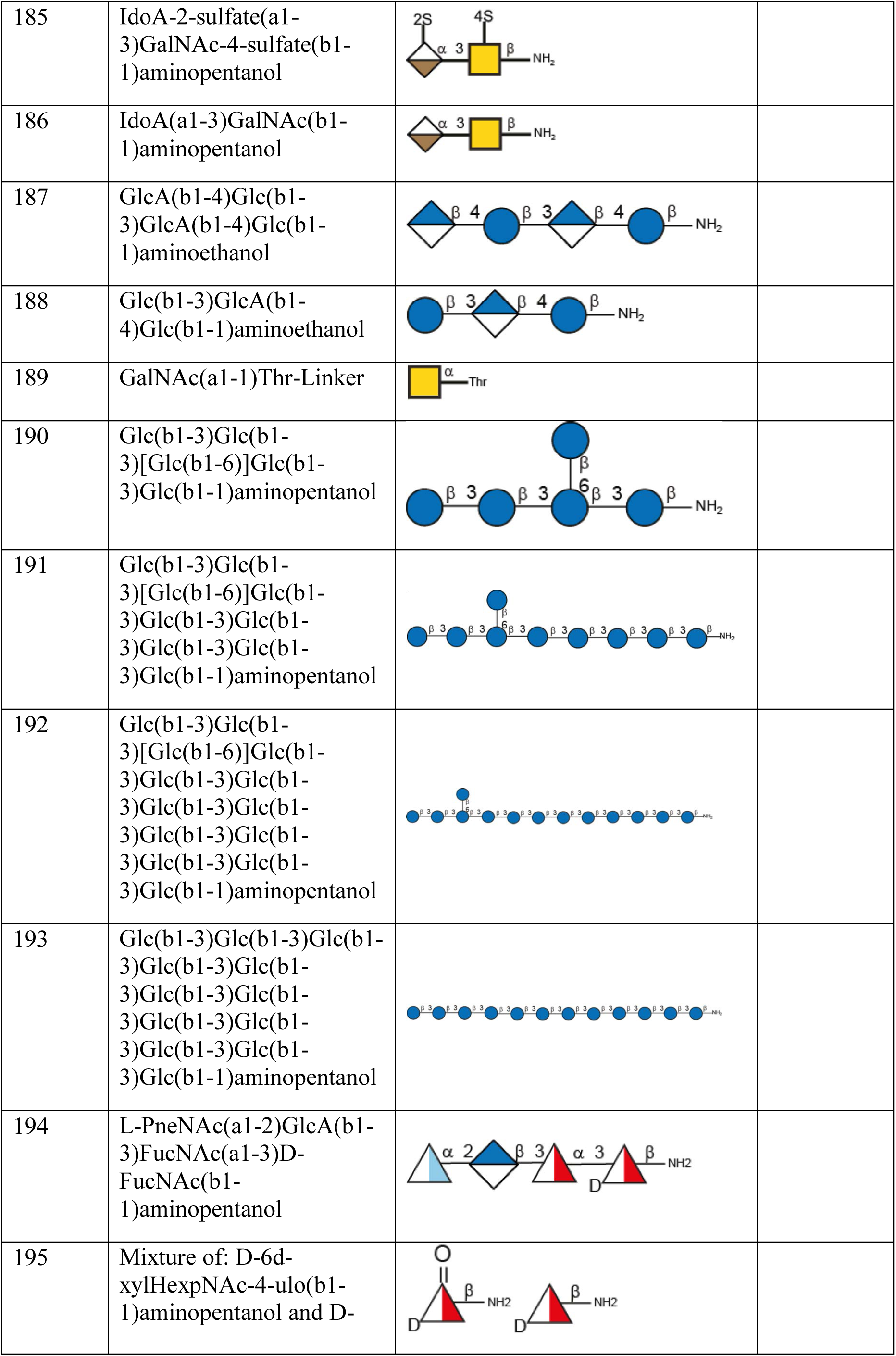
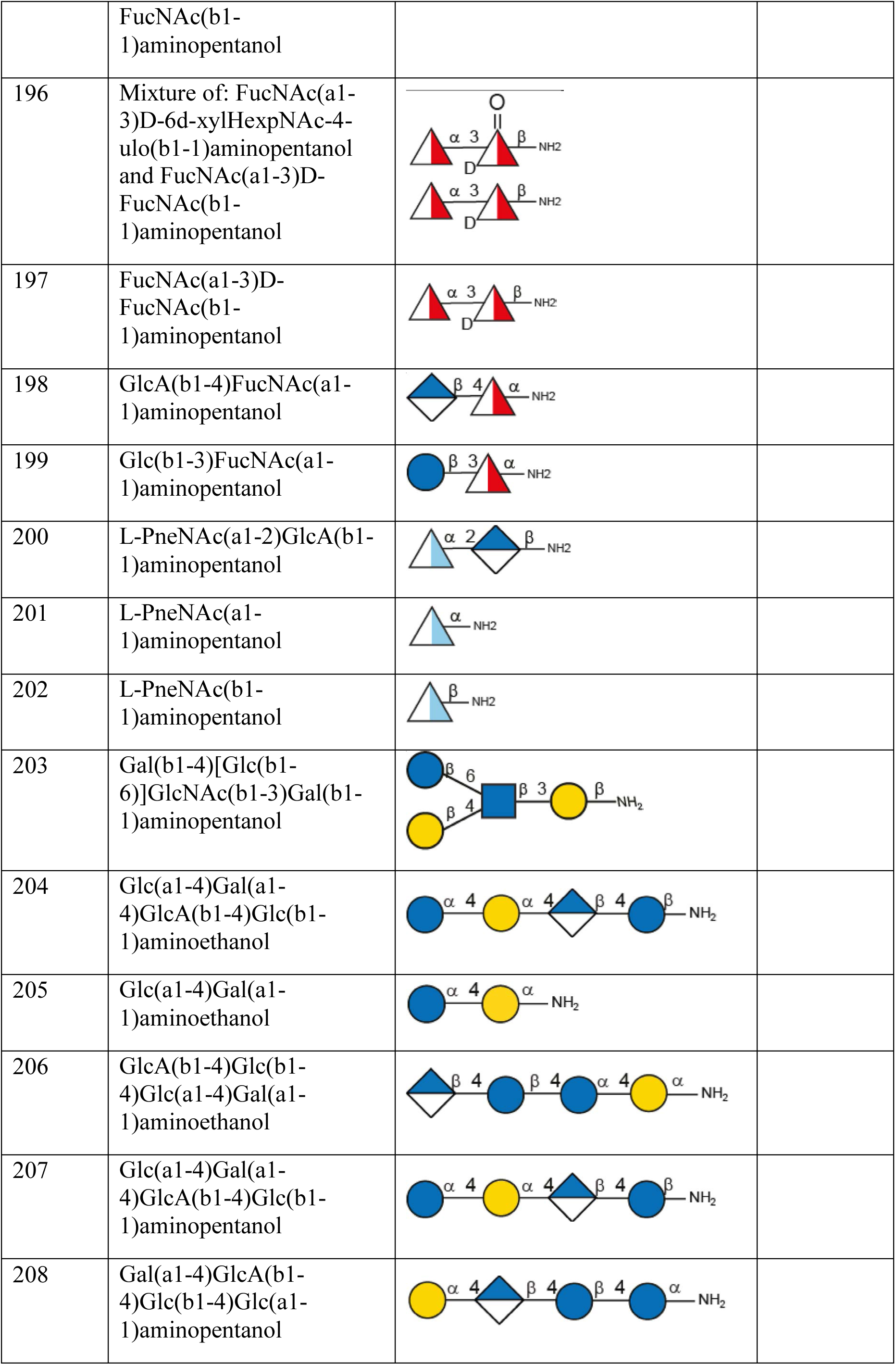
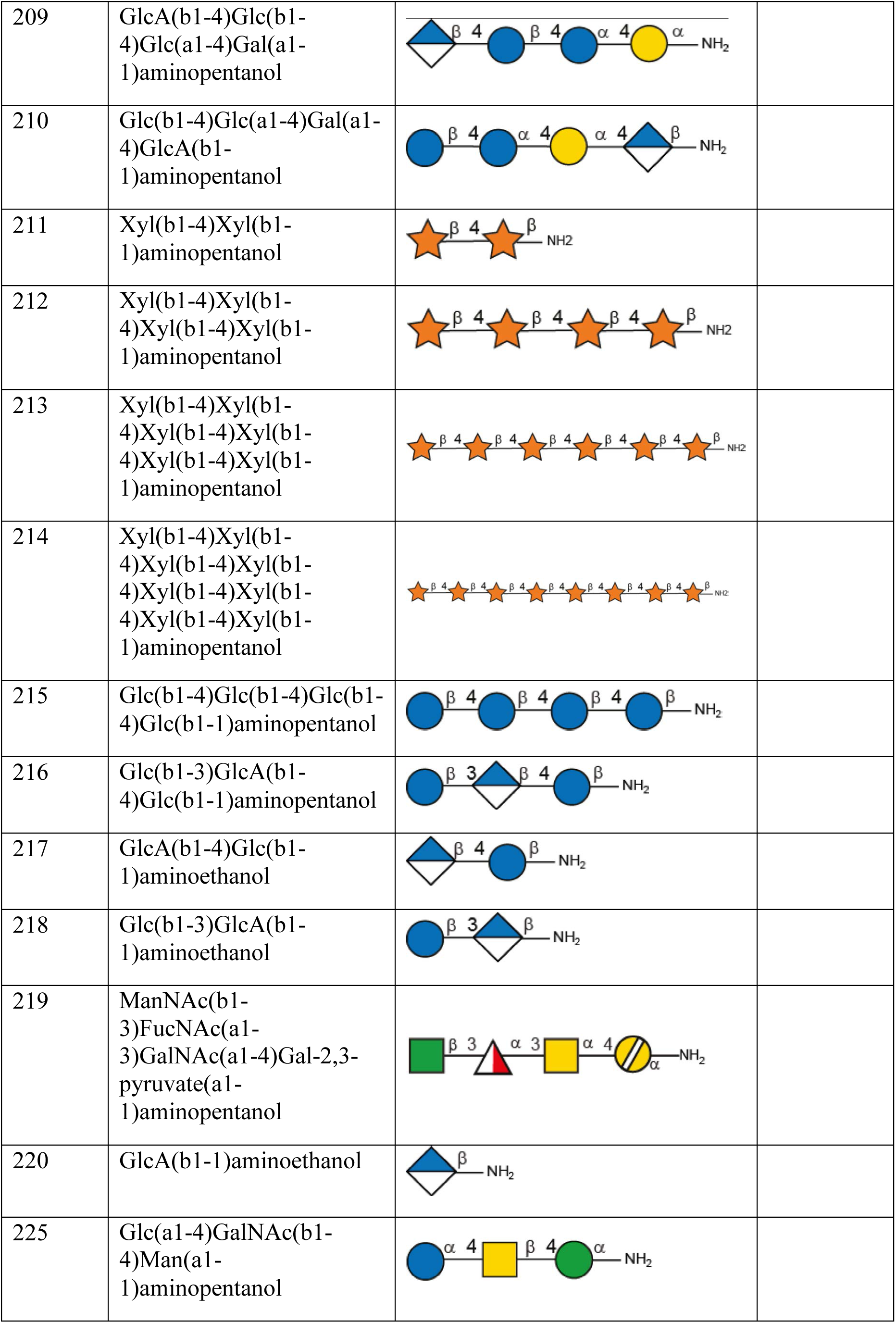
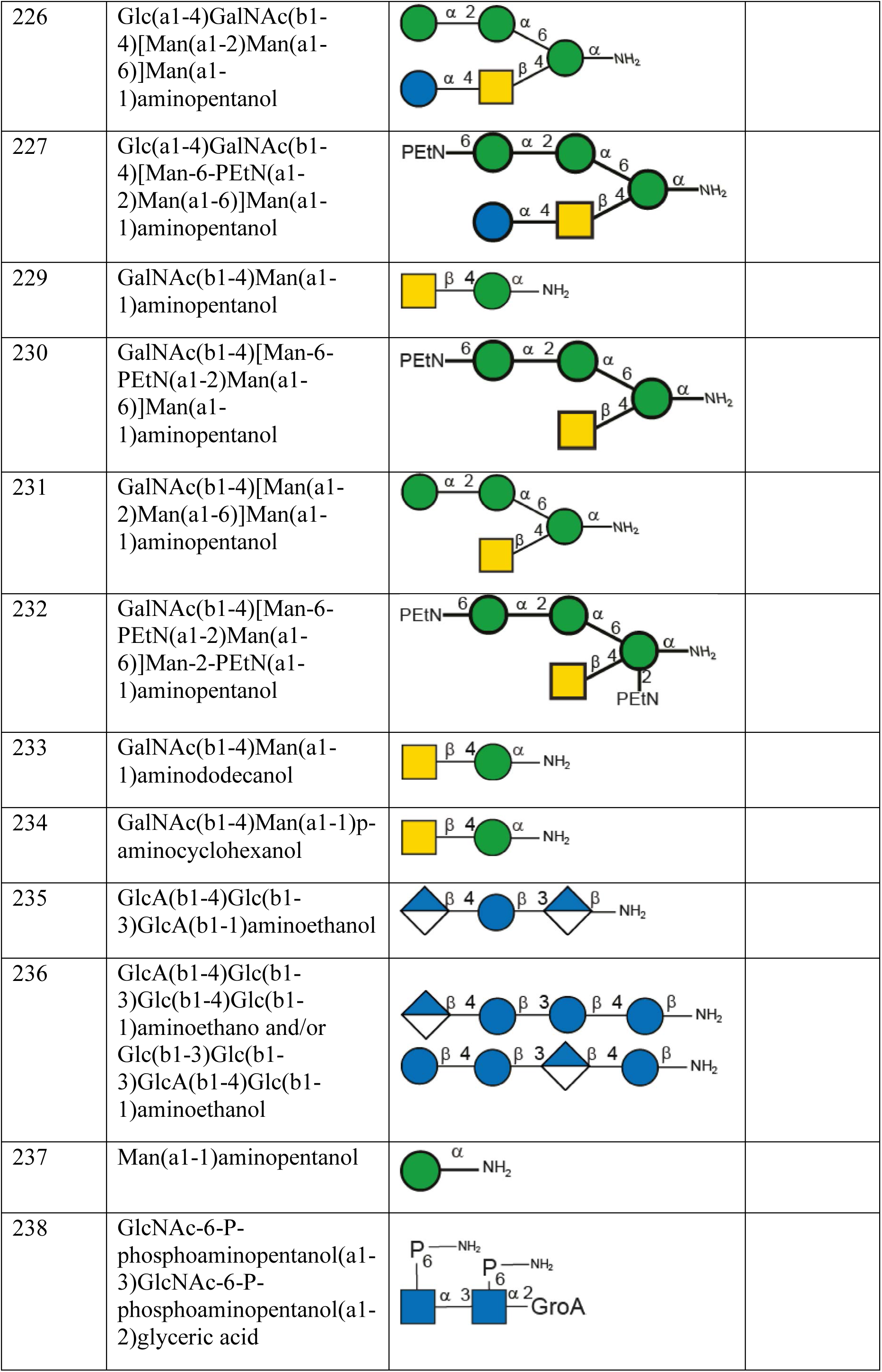
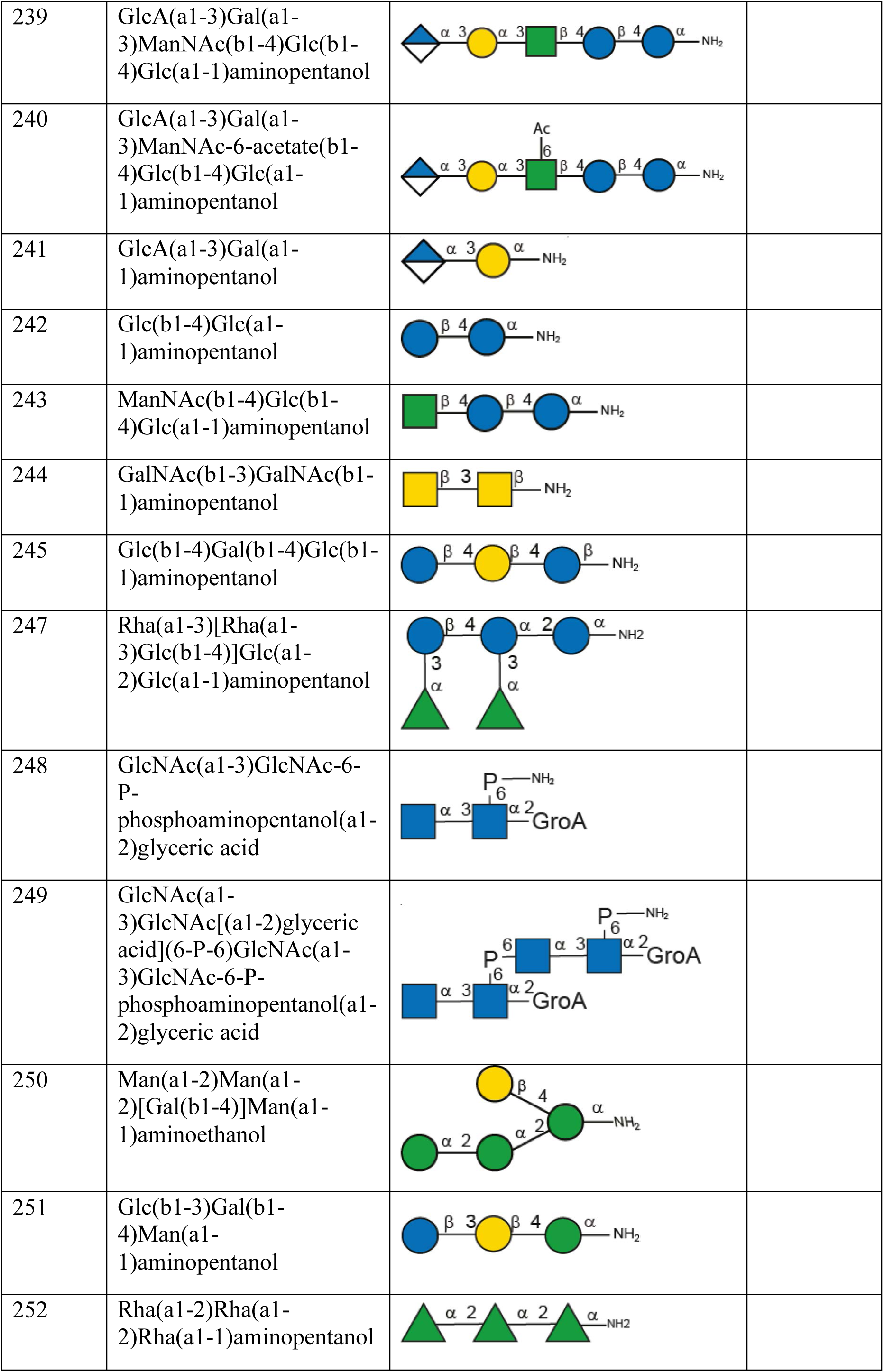
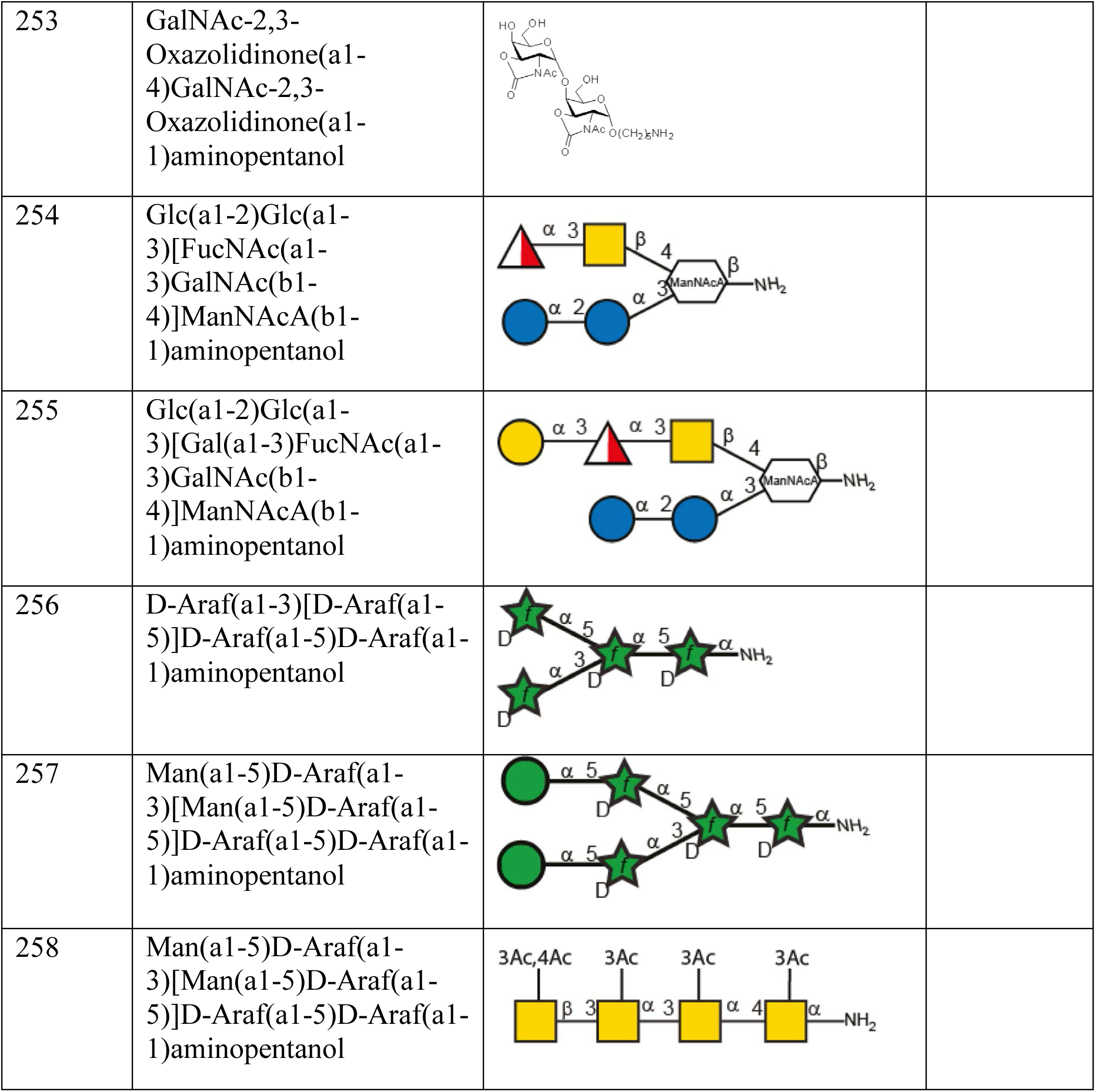
Overview of all glycans on the microarray in Figure 2a. For additional information see reference no. 40.

| Glycan ID | Name | Structure | Compound no. Fig. 2a |
| --- | --- | --- | --- |
| 5 | Neu5Ac(a2-6)Gal(b1-4)GlcNAc(b1-3)Gal(b1-4)Glc(b1-1)aminohexanol |  |  |
| 6 | Neu5Ac(a2-3)Gal(b1-3)GlcNAc(b1-3)Gal(b1-4)Glc(b1-1)aminohexanol |  |  |
| 7 | Fuc(a1-3)[Neu5Ac(a2-3)Gal(b1-4)]GlcNAc(b1-3)Gal(b1-4)Glc(b1-1)aminohexanol |  | 4 |
| 8 | Neu5Ac(a2-6)Gal(b1-4)Glc(b1-1)aminohexanol |  |  |
| 9 | Neu5Ac(a2-3)Gal(b1-4)Glc(b1-1)aminohexanol |  |  |
| 10 | Neu5Ac(a2-6)Gal(b1-4)GlcNAc-6-sulfate(b1-1)aminohexanol |  |  |
| 11 | Gal(b1-4)Glc(b1-1)aminohexanol |  |  |
| 12 | Gal(b1-4)GlcNAc-6-sulfate(b1-1)aminohexanol |  |  |
| 69 | D-Araf(a1-5)D-Araf(a1-1)aminopentanol |  |  |
| 70 | D-Araf(a1-5)D-Araf(a1-3)[D-Araf(a1-5)D-Araf(a1-5)]D-Araf(a1-5)D-Araf(a1-1)aminopentanol |  |  |
| 71 | D-Araf(a1-3)[D-Araf(a1-5)]D-Araf(a1-1)aminopentanol |  |  |
| 72 | D-Araf(a1-5)D-Araf(a1-5)D-Araf(a1-5)D-Araf(a1-5)D-Araf(a1-5)D-Araf(a1-5)D-Araf(a1-1)aminopentanol |  |  |
|  | 5)D-Araf(a1-5)D-Araf(a1-5)aminopentanol |  |  |
| 73 | Col(a1-3)[Col(a1-6)]Glc(a1-4)Gal(a1-3)GlcNAc(b1-1)aminopentanol |  |  |
| 74 | ManNAc(b1-3)FucNAc(a1-3)GalNAc(a1-4)Gal(a1-1)aminopentanol |  |  |
| 75 | GalNAc(a1-4)Gal(a1-1)aminopentanol |  |  |
| 76 | GalNAc(b1-4)Gal(a1-1)aminopentanol |  |  |
| 77 | FucNAc(a1-3)GalNAc(a1-4)Gal(a1-1)aminopentanol |  |  |
| 78 | FucNAc(b1-3)GalNAc(a1-4)Gal(a1-1)aminopentanol |  |  |
| 80 | GalNAc(b1-1)aminoethanol |  |  |
| 81 | FucNAc(a1-1)aminopentanol |  |  |
| 82 | Man(a1-2)Man(a1-2)[Gal(b1-4)]Man(a1-1)aminopentanol |  |  |
| 83 | Man(a1-2)Man(a1-2)[Gal(b1-4)]Man(a1-1)aminopentanol |  |  |
| 84 | Man(a1-2)Man(a1-2)Man(a1-1)aminopentanol |  |  |
| 85 | Gal(b1-4)Man(a1-1)aminopentanol |  |  |
| 90 | Glc(b1-1)aminoethanol |  |  |
| 91 | GlcNAc(a1-2)Hep(a1-3)Hep(a1-5)Kdo(a2-1)aminopentanol |  |  |
| 92 | Hep(a1-3)Hep(a1-5)Kdo(a2-1)aminopentanol |  |  |
| 93 | Hep(a1-3)Hep(a1-5)[L-Ara4N(b1-8)]Kdo(a2-1)aminopentanol |  |  |
| 94 | Hep(a1-7)Hep(a1-3)Hep(a1-5)Kdo(a2-1)aminopentanol |  |  |
| 95 | Hep(a1-2)Hep(a1-3)Hep(a1-5)Kdo(a2-1)aminopentanol |  |  |
| 96 | Hep(a1-5)Kdo(a2-1)aminopentanol |  |  |
| 97 | Hep(a1-7)Hep(a1-3)Hep(a1-1)aminopentanol |  |  |
| 98 | Kdo(a2-8)Kdo(a2-4)Kdo(a2-1)aminopentanol |  |  |
| 99 | Kdo(a2-1)aminopentanol |  |  |
| 100 | Hep(a1-1)aminopentanol |  |  |
| 101 | Glc(b1-1)aminopentanol |  |  |
| 102 | D-FucNAc(b1-1)aminopentanol |  |  |
| 103 | D-FucNAc(b1-1)aminopentanol |  |  |
| 104 | FucNAc(b1-1)aminopentanol |  |  |
| 105 | FucNAc(b1-1)aminopentanol |  |  |
| 153 | Gal(b1-3)GalNAc(a1-1)aminopentanol |  |  |
| 154 | Fuc(a1-3)[Gal(b1-4)]GlcNAc(b1-1)aminopentanol |  | **11** |
| 155 | Neu5Ac(a2-6)GalNAc(a1-1)aminopentanol |  |  |
| 156 | Gal(b1-4)[Gal(b1-4)Glc(b1-6)]GlcNAc(b1-1)aminopentanol |  |  |
| 157 | Fuc(a1-3)[Fuc(a1-2)Gal(b1-4)]GlcNAc(b1-1)aminopentanol |  | 5 |
| 158 | Gal(b1-3)[Fuc(a1-4)]GlcNAc(b1-1)aminopentanol |  | 12 |
| 159 | Fuc(a1-2)Gal(b1-3)[Fuc(a1-4)]GlcNAc(b1-1)aminopentanol |  | 6 |
| 160 | Gal-2,3-Pyruvate(a1-1)aminopentanol |  |  |
| 161 | Gal(a1-3)Gal(b1-4)Glc(b1-1)aminopentanol |  |  |
| 162 | Gal(a1-3)Gal(b1-4)GlcNAc(b1-1)aminopentanol |  |  |
| 163 | Gal(a1-3)Gal(b1-4)GlcNAc(b1-3)Gal(b1-4)Glc(b1-1)aminopentanol |  |  |
| 164 | Gal(b1-4)GlcNAc(b1-3)Gal(b1-4)Glc(b1-1)aminopentanol |  |  |
| 165 | Fuc(a1-2)Gal(b1-3)GlcNAc(b1-3)Gal(b1-4)Glc(b1-1)aminopentanol |  |  |
| 166 | Gal(b1-3)GlcNAc(b1-3)Gal(b1-4)Glc(b1-1)aminopentanol |  |  |
| 167 | Rha(a1-1)aminopentanol |  |  |
| 168 | Rha(a1-3)Glc(b1-1)aminopentanol |  |  |
| 169 | Glc(a1-2)Glc(a1-1)aminopentanol |  |  |
| 170 | Glc(b1-4)Glc(a1-2)Glc(a1-1)aminopentanol |  |  |
| 171 | Rha(a1-3)Glc(b1-4)Glc(a1-1)aminopentanol |  |  |
| 172 | Gal(b1-3)GalNAc(b1-3)Gal(a1-4)Gal(b1-4)Glc(b1-1)aminopentanol |  |  |
| 173 | Neu5Ac(a2-8)Neu5Ac(a2-3)[GalNAc(b1-4)]Gal(b1-4)Glc(b1-1)aminopentanol |  |  |
| 174 | Gal(a1-4)Gal(b1-4)Glc(b1-1)aminopentanol |  |  |
| 175 | GalNAc(a1-1)AminoLinker2 |  |  |
| 176 | Fuc(a1-3)[Gal(b1-4)]GlcNAc(b1-1)AminoLinker2 |  | 7 |
| 177 | GlcNAc(a1-2)Hep(a1-3)Hep(a1-1)aminopentanol |  |  |
| 178 | Hep(a1-3)Hep(a1-1)aminopentanol |  |  |
| 179 | Gal(b1-4)Glc(b1-1)aminopentanol |  |  |
| 180 | GalNAc(b1-4)Gal(b1-4)Glc(b1-1)aminopentanol |  |  |
| 181 | Neu5Ac(a2-3)Gal(b1-4)Glc(b1-1)aminopentanol |  |  |
| 182 | GalNAc-4-sulfate(b1-1)aminopentanol |  |  |
| 183 | IdoA-2,4-disulfate(a1-1)aminopentanol |  |  |
| 184 | IdoA(a1-3)GalNAc-4-sulfate(b1-1)aminopentanol |  |  |
| 185 | IdoA-2-sulfate(a1-3)GalNAc-4-sulfate(b1-1)aminopentanol |  |  |
| 186 | IdoA(a1-3)GalNAc(b1-1)aminopentanol |  |  |
| 187 | GlcA(b1-4)Glc(b1-3)GlcA(b1-4)Glc(b1-1)aminoethanol |  |  |
| 188 | Glc(b1-3)GlcA(b1-4)Glc(b1-1)aminoethanol |  |  |
| 189 | GalNAc(a1-1)Thr-Linker |  |  |
| 190 | Glc(b1-3)Glc(b1-3)[Glc(b1-6)]Glc(b1-3)Glc(b1-1)aminopentanol |  |  |
| 191 | Glc(b1-3)Glc(b1-3)[Glc(b1-6)]Glc(b1-3)Glc(b1-3)Glc(b1-3)Glc(b1-3)Glc(b1-1)aminopentanol |  |  |
| 192 | Glc(b1-3)Glc(b1-3)[Glc(b1-6)]Glc(b1-3)Glc(b1-3)Glc(b1-3)Glc(b1-3)Glc(b1-3)Glc(b1-3)Glc(b1-3)Glc(b1-1)aminopentanol |  |  |
| 193 | Glc(b1-3)Glc(b1-3)Glc(b1-3)Glc(b1-3)Glc(b1-3)Glc(b1-3)Glc(b1-3)Glc(b1-3)Glc(b1-3)Glc(b1-3)Glc(b1-1)aminopentanol |  |  |
| 194 | L-PneNAc(a1-2)GlcA(b1-3)FucNAc(a1-3)D-FucNAc(b1-1)aminopentanol |  |  |
| 195 | Mixture of: D-6d-xylHexpNAc-4-ulo(b1-1)aminopentanol and D- |  |  |
|  | FucNAc(b1-1)aminopentanol |  |  |
| 196 | Mixture of: FucNAc(a1-3)D-6d-xylHexpNAc-4-ulo(b1-1)aminopentanol and FucNAc(a1-3)D-FucNAc(b1-1)aminopentanol |  |  |
| 197 | FucNAc(a1-3)D-FucNAc(b1-1)aminopentanol |  |  |
| 198 | GlcA(b1-4)FucNAc(a1-1)aminopentanol |  |  |
| 199 | Glc(b1-3)FucNAc(a1-1)aminopentanol |  |  |
| 200 | L-PneNAc(a1-2)GlcA(b1-1)aminopentanol |  |  |
| 201 | L-PneNAc(a1-1)aminopentanol |  |  |
| 202 | L-PneNAc(b1-1)aminopentanol |  |  |
| 203 | Gal(b1-4)[Glc(b1-6)]GlcNAc(b1-3)Gal(b1-1)aminopentanol |  |  |
| 204 | Glc(a1-4)Gal(a1-4)GlcA(b1-4)Glc(b1-1)aminoethanol |  |  |
| 205 | Glc(a1-4)Gal(a1-1)aminoethanol |  |  |
| 206 | GlcA(b1-4)Glc(b1-4)Glc(a1-4)Gal(a1-1)aminoethanol |  |  |
| 207 | Glc(a1-4)Gal(a1-4)GlcA(b1-4)Glc(b1-1)aminopentanol |  |  |
| 208 | Gal(a1-4)GlcA(b1-4)Glc(b1-4)Glc(a1-1)aminopentanol |  |  |
| 209 | GlcA(b1-4)Glc(b1-4)Glc(a1-4)Gal(a1-1)aminopentanol |  |  |
| 210 | Glc(b1-4)Glc(a1-4)Gal(a1-4)GlcA(b1-1)aminopentanol |  |  |
| 211 | Xyl(b1-4)Xyl(b1-1)aminopentanol |  |  |
| 212 | Xyl(b1-4)Xyl(b1-4)Xyl(b1-4)Xyl(b1-1)aminopentanol |  |  |
| 213 | Xyl(b1-4)Xyl(b1-4)Xyl(b1-4)Xyl(b1-4)Xyl(b1-4)Xyl(b1-1)aminopentanol |  |  |
| 214 | Xyl(b1-4)Xyl(b1-4)Xyl(b1-4)Xyl(b1-4)Xyl(b1-4)Xyl(b1-4)Xyl(b1-4)Xyl(b1-1)aminopentanol |  |  |
| 215 | Glc(b1-4)Glc(b1-4)Glc(b1-4)Glc(b1-1)aminopentanol |  |  |
| 216 | Glc(b1-3)GlcA(b1-4)Glc(b1-1)aminopentanol |  |  |
| 217 | GlcA(b1-4)Glc(b1-1)aminoethanol |  |  |
| 218 | Glc(b1-3)GlcA(b1-1)aminoethanol |  |  |
| 219 | ManNAc(b1-3)FucNAc(a1-3)GalNAc(a1-4)Gal-2,3-pyruvate(a1-1)aminopentanol |  |  |
| 220 | GlcA(b1-1)aminoethanol |  |  |
| 225 | Glc(a1-4)GalNAc(b1-4)Man(a1-1)aminopentanol |  |  |
| 226 | Glc(a1-4)GalNAc(b1-4)[Man(a1-2)Man(a1-6)]Man(a1-1)aminopentanol |  |  |
| 227 | Glc(a1-4)GalNAc(b1-4)[Man-6-PEtN(a1-2)Man(a1-6)]Man(a1-1)aminopentanol |  |  |
| 229 | GalNAc(b1-4)Man(a1-1)aminopentanol |  |  |
| 230 | GalNAc(b1-4)[Man-6-PEtN(a1-2)Man(a1-6)]Man(a1-1)aminopentanol |  |  |
| 231 | GalNAc(b1-4)[Man(a1-2)Man(a1-6)]Man(a1-1)aminopentanol |  |  |
| 232 | GalNAc(b1-4)[Man-6-PEtN(a1-2)Man(a1-6)]Man-2-PEtN(a1-1)aminopentanol |  |  |
| 233 | GalNAc(b1-4)Man(a1-1)aminododecanol |  |  |
| 234 | GalNAc(b1-4)Man(a1-1)p-aminocyclohexanol |  |  |
| 235 | GlcA(b1-4)Glc(b1-3)GlcA(b1-1)aminoethanol |  |  |
| 236 | GlcA(b1-4)Glc(b1-3)Glc(b1-4)Glc(b1-1)aminoethano and/or Glc(b1-3)Glc(b1-3)GlcA(b1-4)Glc(b1-1)aminoethanol |  |  |
| 237 | Man(a1-1)aminopentanol |  |  |
| 238 | GlcNAc-6-P-phosphoaminopentanol(a1-3)GlcNAc-6-P-phosphoaminopentanol(a1-2)glyceric acid |  |  |
| 239 | GlcA(a1-3)Gal(a1-3)ManNAc(b1-4)Glc(b1-4)Glc(a1-1)aminopentanol |  |  |
| 240 | GlcA(a1-3)Gal(a1-3)ManNAc-6-acetate(b1-4)Glc(b1-4)Glc(a1-1)aminopentanol |  |  |
| 241 | GlcA(a1-3)Gal(a1-1)aminopentanol |  |  |
| 242 | Glc(b1-4)Glc(a1-1)aminopentanol |  |  |
| 243 | ManNAc(b1-4)Glc(b1-4)Glc(a1-1)aminopentanol |  |  |
| 244 | GalNAc(b1-3)GalNAc(b1-1)aminopentanol |  |  |
| 245 | Glc(b1-4)Gal(b1-4)Glc(b1-1)aminopentanol |  |  |
| 247 | Rha(a1-3)[Rha(a1-3)Glc(b1-4)]Glc(a1-2)Glc(a1-1)aminopentanol |  |  |
| 248 | GlcNAc(a1-3)GlcNAc-6-P-phosphoaminopentanol(a1-2)glyceric acid |  |  |
| 249 | GlcNAc(a1-3)GlcNAc[(a1-2)glyceric acid](6-P-6)GlcNAc(a1-3)GlcNAc-6-P-phosphoaminopentanol(a1-2)glyceric acid |  |  |
| 250 | Man(a1-2)Man(a1-2)[Gal(b1-4)]Man(a1-1)aminoethanol |  |  |
| 251 | Glc(b1-3)Gal(b1-4)Man(a1-1)aminopentanol |  |  |
| 252 | Rha(a1-2)Rha(a1-2)Rha(a1-1)aminopentanol |  |  |
| 253 | GalNAc-2,3-Oxazolidinone(a1-4)GalNAc-2,3-Oxazolidinone(a1-1)aminopentanol |  |  |
| 254 | Glc(a1-2)Glc(a1-3)[FucNAc(a1-3)GalNAc(b1-4)]ManNAcA(b1-1)aminopentanol |  |  |
| 255 | Glc(a1-2)Glc(a1-3)[Gal(a1-3)FucNAc(a1-3)GalNAc(b1-4)]ManNAcA(b1-1)aminopentanol |  |  |
| 256 | D-Araf(a1-3)[D-Araf(a1-5)]D-Araf(a1-5)D-Araf(a1-1)aminopentanol |  |  |
| 257 | Man(a1-5)D-Araf(a1-3)[Man(a1-5)D-Araf(a1-5)]D-Araf(a1-5)D-Araf(a1-1)aminopentanol |  |  |
| 258 | Man(a1-5)D-Araf(a1-3)[Man(a1-5)D-Araf(a1-5)]D-Araf(a1-5)D-Araf(a1-1)aminopentanol |  |  |

**Supplementary Table 3:**
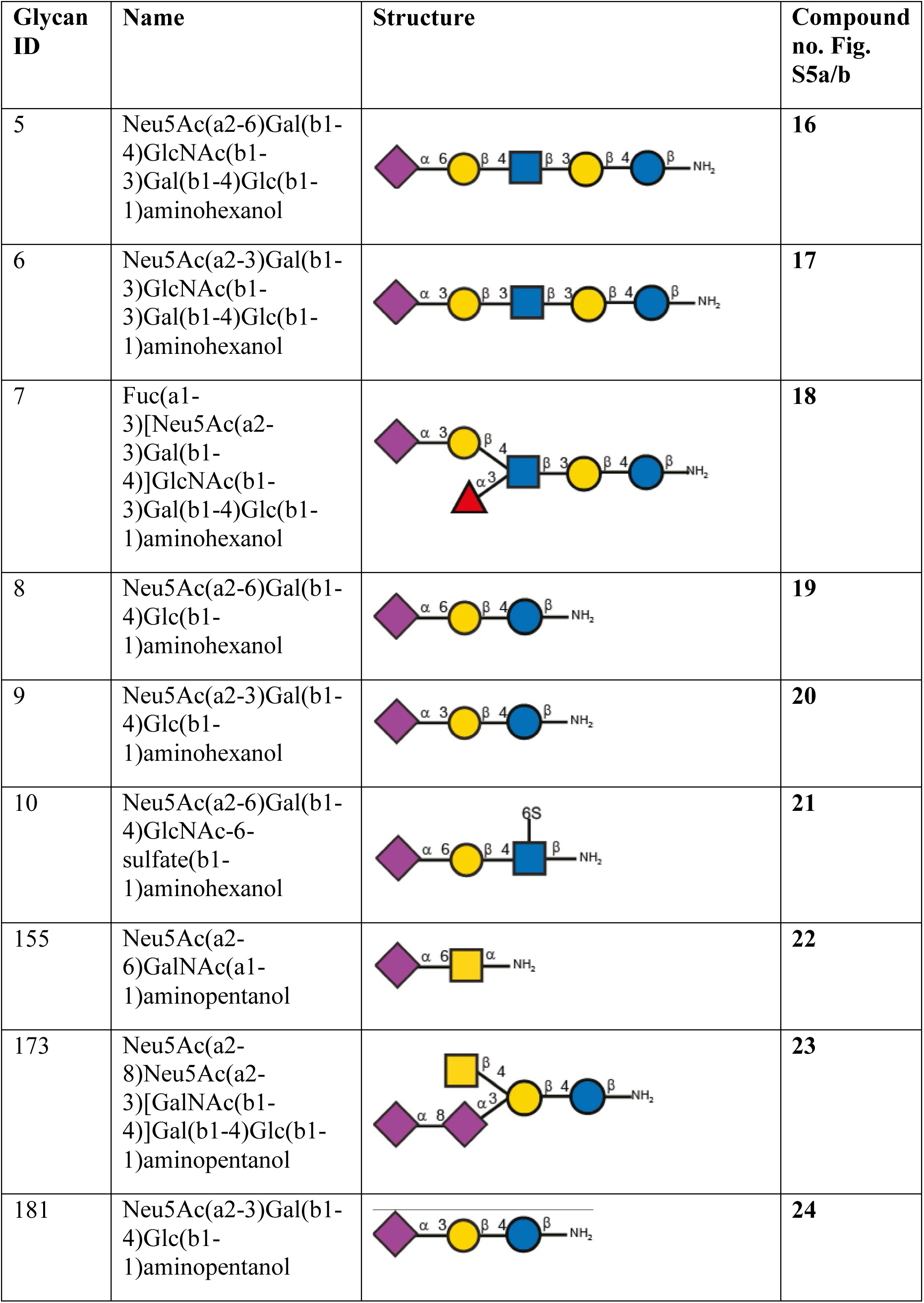
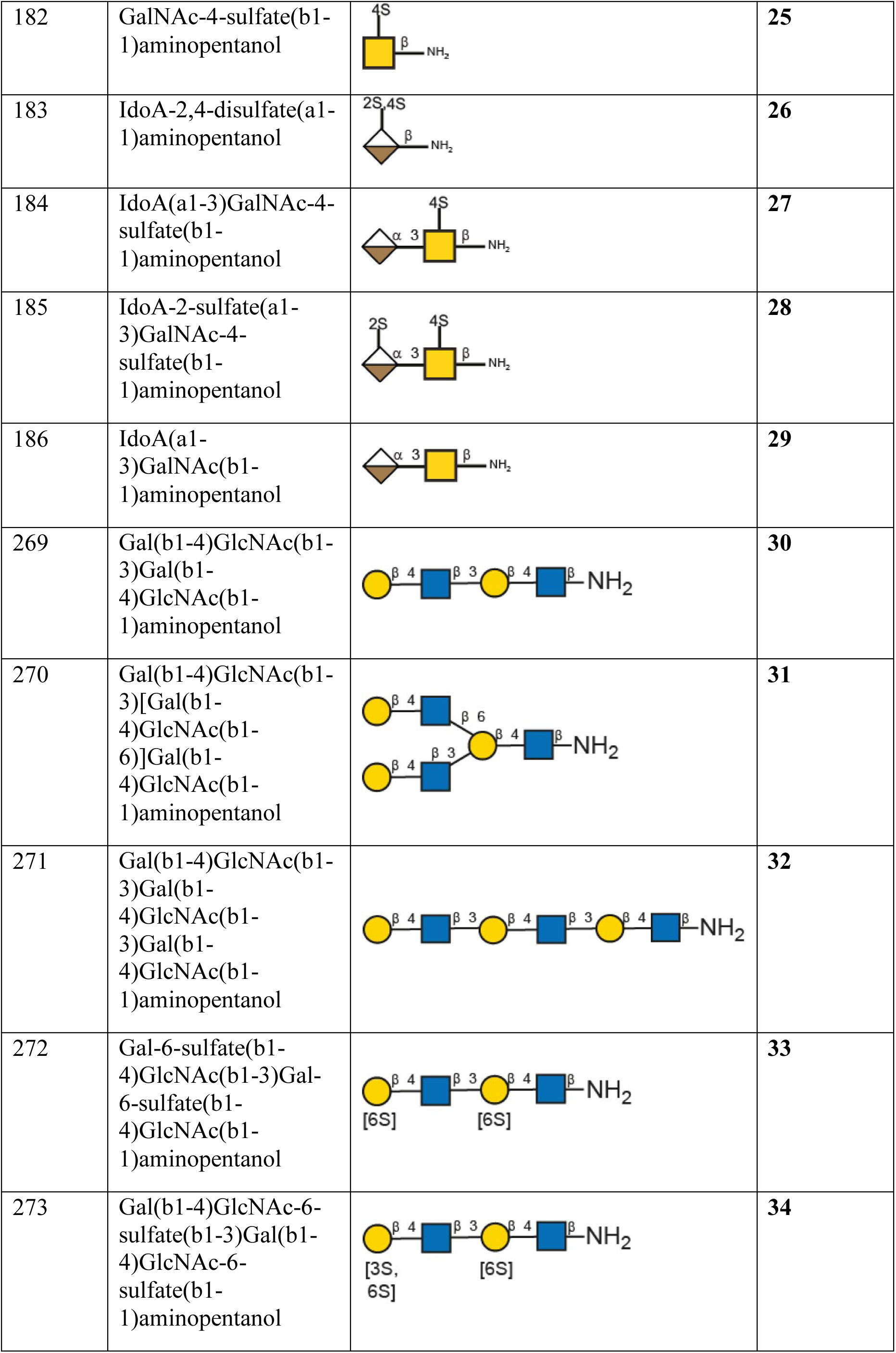
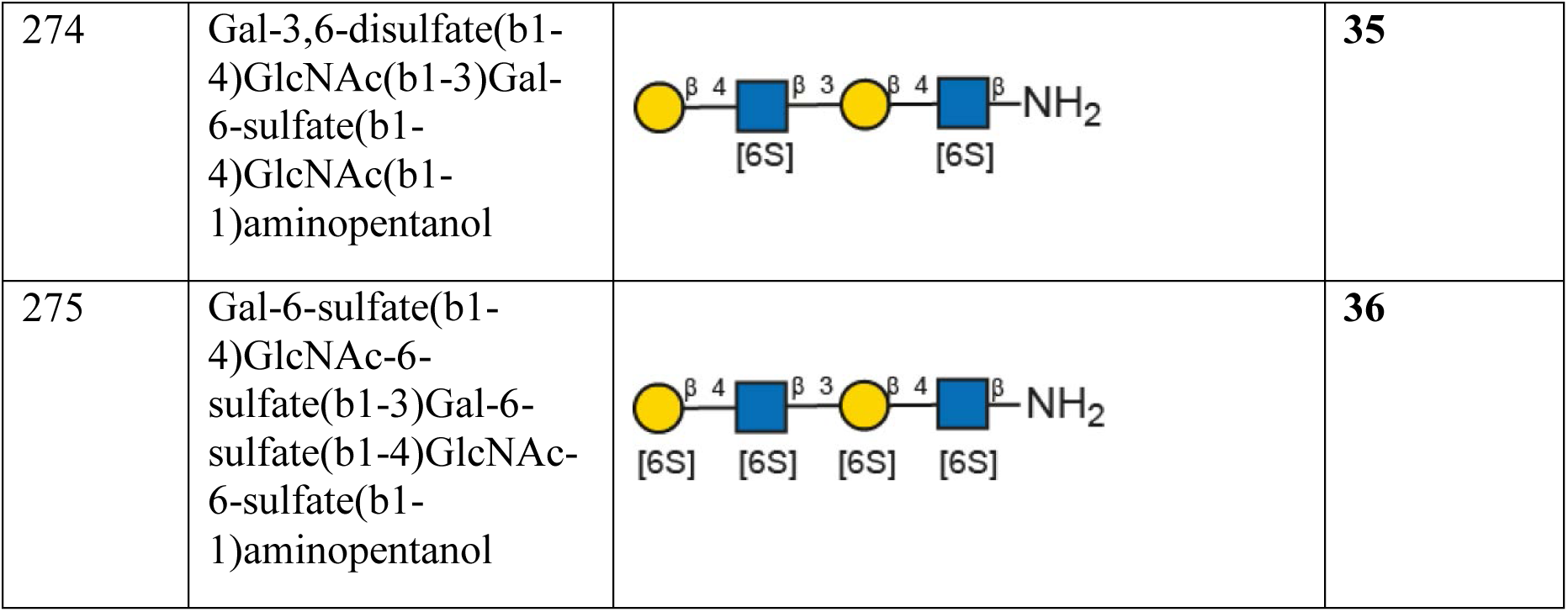
Overview of all glycans on the microarrays in Supplementary Figure 5a,b. For additional information see reference no. 40.

| Glycan ID | Name | Structure | Compound no. Fig. S5a/b |
| --- | --- | --- | --- |
| 5 | Neu5Ac(a2-6)Gal(b1-4)GlcNAc(b1-3)Gal(b1-4)Glc(b1-1)aminohexanol |  | 16 |
| 6 | Neu5Ac(a2-3)Gal(b1-3)GlcNAc(b1-3)Gal(b1-4)Glc(b1-1)aminohexanol |  | 17 |
| 7 | Fuc(a1-3)[Neu5Ac(a2-3)Gal(b1-4)]GlcNAc(b1-3)Gal(b1-4)Glc(b1-1)aminohexanol |  | 18 |
| 8 | Neu5Ac(a2-6)Gal(b1-4)Glc(b1-1)aminohexanol |  | 19 |
| 9 | Neu5Ac(a2-3)Gal(b1-4)Glc(b1-1)aminohexanol |  | 20 |
| 10 | Neu5Ac(a2-6)Gal(b1-4)GlcNAc-6-sulfate(b1-1)aminohexanol |  | 21 |
| 155 | Neu5Ac(a2-6)GalNAc(a1-1)aminopentanol |  | 22 |
| 173 | Neu5Ac(a2-8)Neu5Ac(a2-3)[GalNAc(b1-4)]Gal(b1-4)Glc(b1-1)aminopentanol |  | 23 |
| 181 | Neu5Ac(a2-3)Gal(b1-4)Glc(b1-1)aminopentanol |  | 24 |
| 182 | GalNAc-4-sulfate(b1-1)aminopentanol |  | 25 |
| 183 | IdoA-2,4-disulfate(a1-1)aminopentanol |  | 26 |
| 184 | IdoA(a1-3)GalNAc-4-sulfate(b1-1)aminopentanol |  | 27 |
| 185 | IdoA-2-sulfate(a1-3)GalNAc-4-sulfate(b1-1)aminopentanol |  | 28 |
| 186 | IdoA(a1-3)GalNAc(b1-1)aminopentanol |  | 29 |
| 269 | Gal(b1-4)GlcNAc(b1-3)Gal(b1-4)GlcNAc(b1-1)aminopentanol |  | 30 |
| 270 | Gal(b1-4)GlcNAc(b1-3)[Gal(b1-4)GlcNAc(b1-6)]Gal(b1-4)GlcNAc(b1-1)aminopentanol |  | 31 |
| 271 | Gal(b1-4)GlcNAc(b1-3)Gal(b1-4)GlcNAc(b1-3)Gal(b1-4)GlcNAc(b1-1)aminopentanol |  | 32 |
| 272 | Gal-6-sulfate(b1-4)GlcNAc(b1-3)Gal-6-sulfate(b1-4)GlcNAc(b1-1)aminopentanol |  | 33 |
| 273 | Gal(b1-4)GlcNAc-6-sulfate(b1-3)Gal(b1-4)GlcNAc-6-sulfate(b1-1)aminopentanol |  | 34 |
| 274 | Gal-3,6-disulfate(b1-4)GlcNAc(b1-3)Gal-6-sulfate(b1-4)GlcNAc(b1-1)aminopentanol |  | **35** |
| 275 | Gal-6-sulfate(b1-4)GlcNAc-6-sulfate(b1-3)Gal-6-sulfate(b1-4)GlcNAc-6-sulfate(b1-1)aminopentanol |  | **36** |

